# Structural maturation of the matrix lattice is not required for HIV-1 particle infectivity

**DOI:** 10.1101/2024.12.22.629981

**Authors:** Long Chen, Yuta Hikichi, Juan S. Rey, Caner Akil, Yanan Zhu, Hana Veler, Yao Shen, Juan R. Perilla, Eric O. Freed, Peijun Zhang

## Abstract

HIV-1 assembly is initiated by the binding of Gag polyproteins to the inner leaflet of the plasma membrane, mediated by the myristylated matrix (MA) domain of Gag. Subsequent to membrane binding, Gag oligomerizes and buds as an immature, non-infectious virus particle, which, upon cleavage of the Gag precursor by the viral protease, transforms into a mature, infectious virion. During maturation, the MA lattice underlying the viral membrane undergoes a structural rearrangement and the newly released capsid (CA) protein forms a mature capsid that encloses the viral genome. While it is well established that formation of the mature capsid is essential to particle infectivity, the functional role of MA structural maturation remains unclear. Here, we examine MA maturation of an MA triple mutant, L20K/E73K/A82T, which exhibits distinct biochemical behaviours. The L20K/E73K/A82T mutant is a revertant derived by propagating the L20K mutant, which exhibits reduced infectivity and increased association of the Gag polyprotein with membranes. L20K/E73K/A82T replicates similarly to wild type but retains the increased Gag membrane binding properties of L20K. L20K/E73K/A82T MA also sediments to high-density fractions in sucrose gradients after detergent treatment under conditions that fully solubilize WT MA, suggesting enhanced MA-MA interactions. Cryo-electron tomography with subtomogram averaging reveals that the immature MA lattice of L20K/E73K/A82T closely resembles the wild type. However, mature virions of the triple mutant lack a detectable MA lattice, in stark contrast to both the wild type and L20K mutant. All-atom molecular dynamics simulations suggest that this absence results from destabilized inter-trimer interactions in the mature L20K/E73K/A82T MA. Furthermore, introducing additional mutations designed to disrupt the mature MA lattice does not impair particle infectivity. These findings suggest that an ordered, membrane-associated mature MA lattice is not essential for HIV-1 infectivity, providing new insights into the structural plasticity of the matrix during maturation and its functional role in the viral lifecycle.

## Introduction

The HIV-1 Gag polyprotein precursor, Pr55Gag, includes several domains that serve critical functions during the assembly process: the matrix (MA) domain targets Gag to the plasma membrane and promotes the incorporation of the viral envelope (Env) glycoproteins into virions, the capsid (CA) domain drives Gag oligomerization, the nucleocapsid (NC) domain recruits the RNA genome into assembling particles and promotes particle assembly[1], and the p6 domain facilitates the release of newly assembled virions via recruitment of the cellular ESCRT machinery to the site of budding [2–6].

During virus assembly, a primary role of the MA domain is to bring Gag to the inner leaflet (IL) of the plasma membrane via its covalently attached N-terminal myristate group and highly basic region (HBR) [7, 8]. The MA HBR promotes Gag membrane binding by engaging in electrostatic interactions with the IL of the plasma membrane [9]. Phosphatidylinositol-4,5-bisphosphate [PI(4,5)P_2_] is essential for anchoring Gag to the plasma membrane [10, 11] and maintaining its membrane association [12]. Previous research has demonstrated that PI(4,5)P_2_ directly interacts with HIV-1 MA, inducing a conformational shift that triggers exposure of the myristate [13]. Furthermore, headgroups of phosphatidylserine (PS), phosphatidylethanolamine (PE), and phosphatidylcholine (PC) interact with basic MA residues facing the membrane [14]. The HIV-1 MA protein forms trimers that assemble into a hexameric lattice with central cavities at points of sixfold symmetry on PS-and PI(4,5)P_2_-enriched membranes [15–17]. During HIV-1 maturation, the MA lattice undergoes structural rearrangement, with mature virions displaying a more regular hexameric lattice and smaller central hexameric apertures compared to their immature counterparts [18]. Previous studies have shown that the MA domain of HIV-1 Gag is crucial for Env incorporation into virions [19–24]. Mutations that impair MA trimerization block Env incorporation [23, 24] and can be rescued by second-site mutations in MA that restore MA trimer formation, Env incorporation, and particle infectivity [25]. Notably, many mutations in MA that are defective in Env incorporation cluster around the peripheral tips of the MA trimer near the six-fold cavity [26]; their compensatory mutations, however, arise at the MA trimer interface [23], further implicating the MA trimer as a key structural element in Env incorporation [23–25]. Notably, mutations and deletions in MA that impair Env incorporation can also be effectively addressed by truncating the long gp41 CT [21, 27, 28]. Importantly, Env in immature virions is fusion incompetent. Fusogenicity can be activated by removing the long cytoplasmic tail (CT) of the transmembrane Env glycoprotein gp41 or upon cleavage of Gag by the viral protease (PR) [29, 30]. Gag cleavage by PR induces clustering of Env trimers on the virion that depends on the gp41 CT [31]. These findings raise the possibility that the structural rearrangement of the MA lattice during virion maturation may modulate Env function, which is critical for successful fusion and entry in the next round of infection.

To understand how the conformational rearrangement of the MA lattice affects viral infectivity, we investigated a previously described revertant of the L20K MA mutant, L20K/E73K/A82T [32]. Leu20 in MA is a highly conserved residue located within the HBR [33]. The L20K mutation increases Gag association with membrane but reduces HIV-1 infectivity and viral DNA synthesis, relative to the wild-type (WT) [34]. Acquisition by L20K of the two additional mutations, E73K and A82T, rescues L20K defects in virus replication in a cell-type dependent manner, without reversing the L20K-imposed increase in Gag membrane binding. Intriguingly, unlike the WT and L20K MA, L20K/E73K/A82T MA is resistant to detergent treatment and co-sediments with the core complex [32]. We previously reported that HIV-1 PR cleaves the CT of murine leukemia virus (MLV) Env when it is incorporated into HIV-1 virions and that the L20K/E73K/A82T mutant exhibits a block in this PR-mediated MLV Env cleavage [35]. These intriguing features of the L20K/E73K/A82T mutant stimulated further investigations of its MA lattice during maturation within native virions using cryo-electron tomography (cryoET), in conjunction with computer simulations, biochemical and virological studies.

## Results

### L20K/E73K/A82T MA forms detergent-resistant oligomers

Previous studies demonstrated that the L20K mutation, which increases membrane binding of Gag, causes defects in viral replication, single-cycle infectivity, and post-entry events compared with WT HIV-1 [34, 36]. By propagating the L20K mutant, we identified compensatory second-site mutations in MA (E73K and A82T) that partially rescued replication defects caused by L20K without reversing the increased Gag membrane binding exhibited by L20K [32]. Western blotting of virion-associated proteins confirmed that the L20K and L20K/E73K/A82T mutations did not affect the amount of p17 (MA), p24 (CA), Pr55Gag, or gp41 (Env) in the virion fraction (Fig. 1a). However, this analysis revealed that both L20K and L20L/E73K/A82T displayed two additional bands, not observed in WT virions, at approximately 34 and 43 kDa (Fig.1a). To determine the identity of these additional bands, we probed with anti-p17 antibodies (Fig 1b). Consistent with the data in Fig 1a, L20K and L20K/E73K/A82T showed two bands at 34 and 43kDa, suggesting that these bands correspond to a MA dimer and trimer, respectively. The L20K/E73K/A82T mutant exhibits more MA dimer/trimer than L20K (Fig. 1c), suggesting that the compensatory mutations enhance detergent-resistant MA-MA interactions.

**Figure 1.**
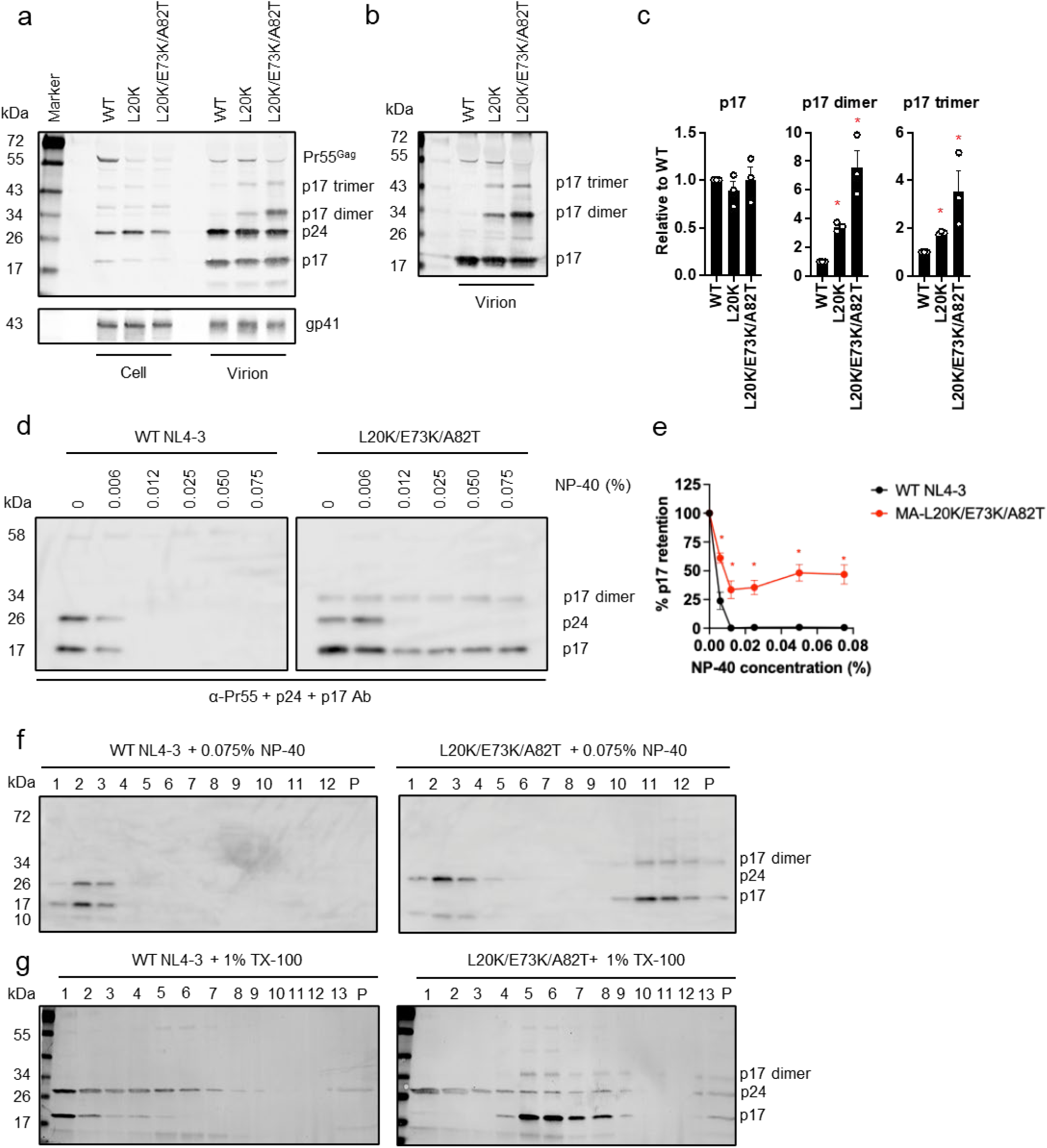
MA-L20K/E73K/A82T exhibits strong MA-MA interactions. a-c) Gag processing, Env expression and incorporation (a) cell and virion-associated proteins derived from 293T cells transfected with WT or MA-mutant pNL4-3 molecular clones were probed with anti-Pr55Gag + p24 (CA) + p17 (MA) Ab and anti-Env (gp41) Ab. (b) virion-associated proteins were probed with anti-p17 (MA) polyclonal Ab. Representative western blots from three independent experiments are shown. (c) Quantification of p17 monomer/dimer/trimer levels from western blots in (b). The data are shown as means ± SEM from three independent experiments with statistical significance indicated (*p < 0.05; Student’s t-test). (d -e) Membrane stripping assay using the indicated concentration of NP-40. (d) Representative western blots from three experiments. (e) Quantification of p17 monomer levels from western blots in (d), indicating the % of p17 retained following detergent treatment relative to the amount present in the absence of detergent. The data are shown as means ± SEM from three independent experiments with statistical significance indicated (*p < 0.05; Student’s t-test). (f) Sedimentation analysis in a 10-70% sucrose gradient after stripping the viral membrane with 0.075% NP-40. Representative western blots from three independent experiments are shown. (g) Sedimentation analysis in 3-60% optiprep gradient after stripping the viral membrane with 1% Triton X-100 (TX-100). Representative western blots from three independent experiments are shown. The western blots of the gradient fractions were analyzed using antibodies against Pr55, p24 (CA), and p17 (MA) for both the WT and mutant MA. P, pellet fraction.

To further examine the stability of the L20K/E73K/A82T MA dimer and trimer, we exposed viruses to a range of concentrations of Nonidet P-40 (NP-40) and analyzed viral proteins in the pelleted fraction (Fig. 1d). While WT NL4-3 CA and MA were completely solubilized by 0.012% NP-40, the L20K/E73K/A82T MA sedimented in the pelleted fraction in the presence of even higher concentrations of NP-40, with the detergent-resistant MA including both monomer and dimer forms (Fig. 1d-e). To examine the density profiles of the detergent-resistant MA monomer and multimers, we performed equilibrium sedimentation analysis after exposure of viruses to 0.075% NP-40. (Fig. 1f). Under conditions that disrupt the capsid, the mutant MA monomer and multimers preferentially co-sedimented with higher-density fractions, suggestive of the formation of a protein ’skeleton’ induced by enhanced MA-MA interactions (Fig. 1f). To extend this finding, we isolated viral cores using a ‘spin-through’ detergent treatment method with 1% TX-100 [37]. While most of WT MA was stripped from the viral core fraction (fraction 5 – 7) under these conditions, the mutated MA co-sedimented with the viral core (Fig. 1g), suggesting that the MA ‘skeleton’ structure maintained its integrity in the presence of the detergent. Taken together, these results suggest a distinct pattern of MA-MA interactions induced by the L20K/E73K/A82T mutations. As the L20K/E73K/A82T mutant displays near-WT infectivity and replication kinetics in primary peripheral blood mononuclear cells (PBMCs) and certain T-cell lines (e.g. H9 cells) [32], these results indicate that the unusual biochemical properties of this mutant do not compromise particle infectivity. These observations prompted us to carry out structural analysis of the L20K/E73K/A82T mutant.

### CryoET of immature and mature L20K/E73K/A82T VLPs

To understand the unusual properties of the L20K/E73K/A82T MA mutant, we imaged immature and mature virus-like particles (VLPs) of this mutant using cryoET and compared them with WT VLPs. VLPs were produced from CMV-promoter driven GagPol expression vectors expressing either WT PR (for mature particles) or a mutant PR containing the inactivating D25N mutation (for immature VLPs) (Fig. 2a), along with Env-and Tat-expressing plasmids. Raw tomograms of immature VLPs displayed a similar morphology for WT and the L20K/E73K/A82T MA mutant, showing striated CA and MA layers (Fig. 2, b-e, blue and gold arrowheads, respectively), indicating lattice formation. In mature VLPs, while conical CA cores are apparent for both WT and the L20K/E73K/A82T mutant (Fig. 2f, I, blue arrowheads), the mutant MA layers look different from the WT (Fig. 2i-k). The mature WT VLPs display a regular, striated MA lattice just below the IL of the lipid bilayer (Fig. 2f, gold arrowhead) and clear lattice arrays in tomographic slices sectioning along the membrane (Fig. 2h), as previously reported [18]. Interestingly, these features were absent in the mature L20K/E73K/A82T VLPs. No MA lattice was evident underneath the IL of the membrane in mature L20K/E73K/A82T VLPs (Fig. 2i-k). Interestingly, while we observed clustering of Env trimers on the surface of both WT and L20K/E73K/A82T mature VLPs, consistent with the previous fluorescence nanoscopic observations [31], these Env clusters seem to associate with the surface area displaying no apparent MA lattice (Fig. 2l-o).

**Figure 2.**
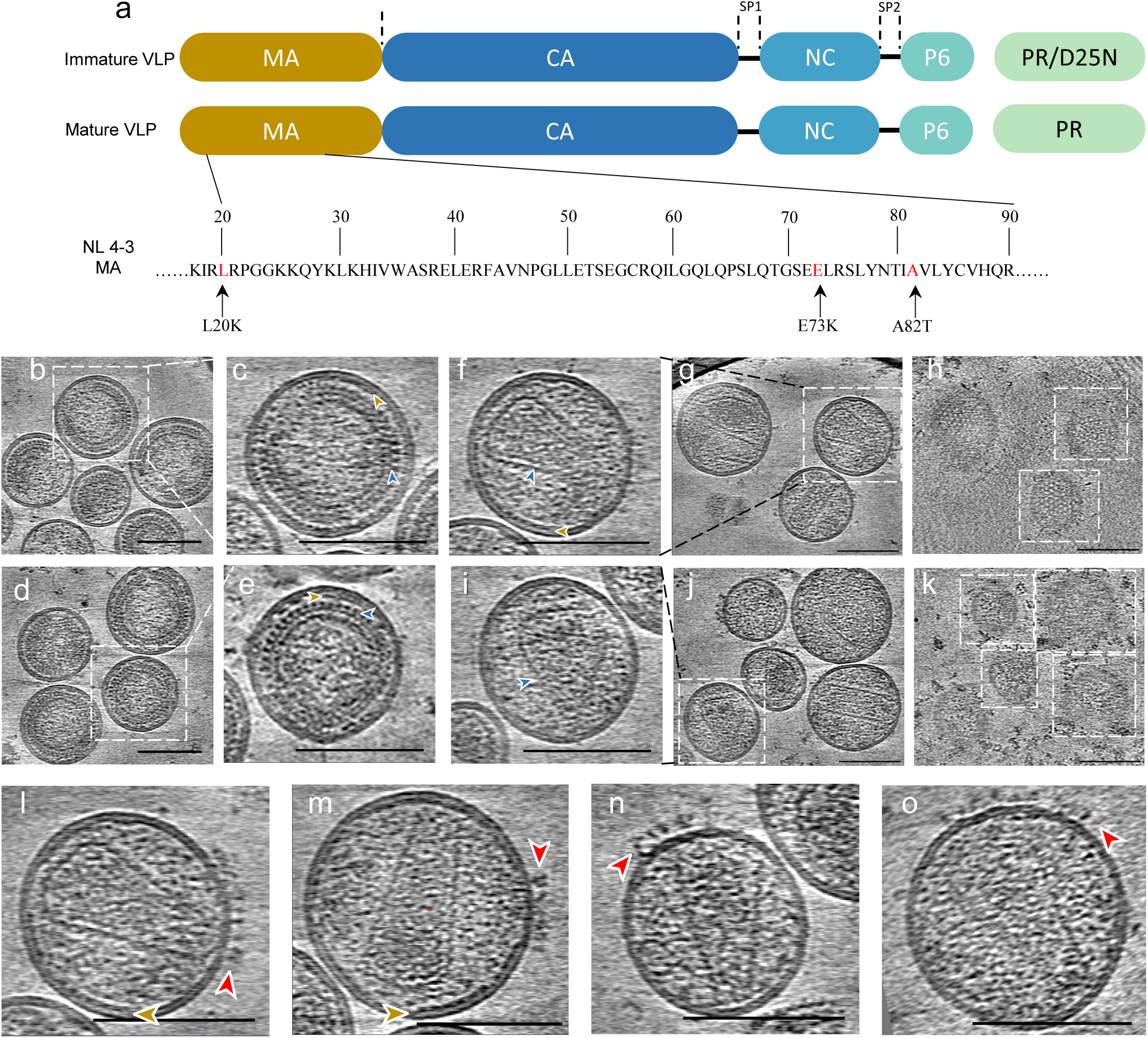
CryoET of immature and mature HIV-1 VLPs of WT and L20K/E73K/A82T mutant. a) Schematic showing WT and L20K/E73K/A82T MA-mutant Gag used to generate the immature and mature VLPs, with proteolytic cleavage sites indicated by vertical dotted black lines. In the immature VLPs, the PR-D25N mutation inactivates the viral PR. The p17 (MA) protein is illustrated with the corresponding amino acid sequence of the WT pNL4-3 MA (residues 17-90). Red indicates amino acid changes in the L20K/E73K/A82T mutant. b-e) CryoET central slices of immature WT (b, c) and mutant (d, e) particles, enhanced by summing 10 neighboring slices. The density appears in black with the MA layer and CA layer highlighted by brown and blue arrowheads, respectively. (f-k) Computational slices through tomographic reconstructions of mature WT (f-h) and L20K/E73K/A82T mutant (i-k) particles. Panels f, g, i, and j show central slices enhanced by summing 10 neighboring slices, displaying side views of conical capsid cores, denoted by blue arrowheads, and MA layers beneath the inner membrane marked by brown arrowheads. The WT shows a regular lattice pattern, while the mutant exhibits disorganized density. Top views (h-k) focus on slices near the inner membrane surface, showing disrupted MA lattice density in the mutant (k) compared to the regular MA lattice in the WT (h). Gold fiducial markers have been removed from the images for clarity. (l-o) The Env-containing tomographic slices from mature WT (l-m) and mutant (n-o) VLPs. Brown arrowheads indicate the MA lattice, and red arrowheads highlight the Env glycoproteins. Scale bar: 100nm.

### Structures of immature L20K/E73K/A82T MA trimer and CA hexamer

We further carried out subtomogram averaging of immature MA and CA layers using emClarity and Relion software packages [38, 39] (Fig. S1) and obtained immature MA trimer maps for both WT and the L20K/E73K/A82T mutant at 8.0 Å and 8.3 Å resolution, and CA hexamer maps at 4.9 Å and 5.2 Å resolution, respectively (Fig. 3, Table S1, Fig. S2). The MA trimer maps were fitted with the molecular model 7OVQ [18] by rigid-body fitting. Comparison of the WT and L20K/E73K/A82T mutant maps revealed a difference at the trimer centre, where the central pore of the MA trimer is filled in the WT but open in the mutant (Fig. 3a-b, Fig. S2c-d, red arrowhead). To evaluate whether the observed difference is statistically significant, we calculated the difference map between the mutant and the WT MA trimers thresholded at 1% false discovery rate, where density gain and loss are colored in blue and red, respectively (Fig. 3c-d). The difference map displayed pairs of density “gain” and “loss” which are largely associated with the mutations (Fig. 3c-d, L20K and A82T). Interestingly, in addition to the significant density loss in the center of the L20K/E73K/A82T MA trimer, there was an unpaired density gain at the C-terminus of L20K/E73K/A82T MA pointing to the CA layer (Fig. 3d, blue arrow), suggesting that this region is likely less flexible than the WT counterpart. Additionally, the electrostatic potential on the surface of MA facing the membrane is also more positive than the WT, due to the L/K and E/K changes in the L20K/E73K/A82T mutant (Fig. 3e-f). Further comparative analysis on the spatial arrangement of MA trimers and the underlying CA hexamers, namely the distance and angle between the nearest neighbors, showed a consistent profile between WT and L20K/E73K/A82T (Fig. 3g-h, Fig. S2g-h), suggesting that the structural organization of the immature MA and CA lattices is highly preserved in the L20K/E73K/A82T mutant VLPs and shows a high degree of resilience to these mutations.

**Figure 3.**
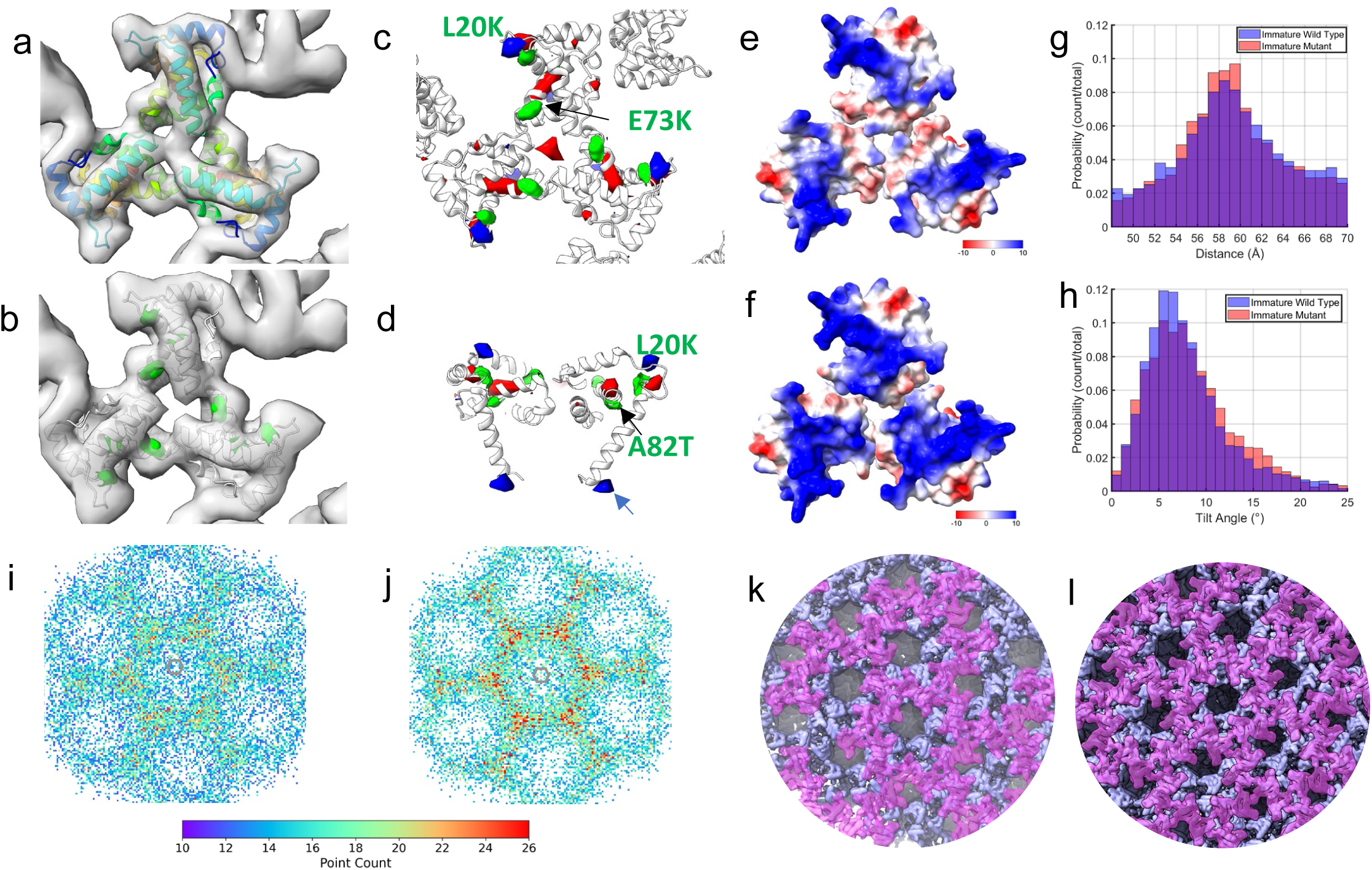
Comparison of immature WT and L20K/E73K/A82T MA mutant. a-b) Subtomogram averaging maps of the MA trimer for the WT (a) and L20K/E73K/A82T mutant (b) are shown as gray isosurfaces, viewed from the top toward the center of the virus. The molecular model of 7OVQ [18] was fitted into both maps as a rigid body. In the WT, the model is colored from blue (N-terminus) to red (C-terminus), while in the mutant, the mutations L20K, E73K, and A82T are highlighted in green. (c-d) The differential density maps, calculated as mutant minus WT, reveal statistically significant changes at a 1% false discovery rate threshold. The green density marks the locations of mutations. Red indicates regions of disappearing density, while blue indicates regions of appearing density. (e-f) Surface electrostatic potential of the MA trimer, derived from Alphafold2-multimer predictions in Colab [108, 109] and molecular dynamics simulations, are displayed for the WT (e) and mutant (f). The maps represent electrostatic charge distribution, with red indicating negatively charged areas and blue indicating positively charged areas. (g-h) Comparative analysis of distances and tilt angles between trimer-trimer pairings in tomographic reconstructions between the WT (blue) and mutant (orange). The MA-MA distance for WT averaged 59.3 Å with a standard deviation of 5.2 Å, while the mutant recorded 59.2 Å with a standard deviation of 5.0 Å. Tilt angles for the WT were 7.6° (SD 3.9°) versus the mutant’s 8.1° (SD 4.2°). (i-l) Radial registration analysis between the MA trimer and CA hexamer. After subtomogram averaging of MA and CA, the reconstructions were generated by mapping back the refined positions and orientations. The MA trimer was then projected onto the CA layer to identify intersection points. A heatmap of these intersection points relative to the CA hexamer shows a denser concentration of MA-CA registration in the mutant (j) compared to the WT (i). Representative reconstruction maps illustrate the spatial positioning of MA-CA interactions for the WT (k) and L20K/E73K/A82T mutant (l). A total of 8000 MA trimers were used for the projection in both the WT and mutant.

### The L20K/E73K/A82T mutations enhance immature MA and CA lattice registration

In immature Gag, the MA domain, which is anchored to the membrane and forms a trimeric lattice, is flexibly linked to the CA domain, which forms a lattice of hexamers below the MA layer. We hypothesize that the MA lattice might be in register with the CA lattice. To visualize both the MA trimer lattice and the CA hexamer lattice, we placed MA trimer and CA hexamer sub-volumes back in the original tomograms (Fig. S3a-b). Subsequent analysis of MA trimer localization from WT and L20K/E73K/A82T mutant VLPs showed that the MA lattices are very similar but are less well ordered than the CA lattice (Fig. S3c-f).

We next investigated whether the linker between MA and CA constrains the relative position of these two domains and whether the MA and CA lattices are in register. To understand the relationship between MA and CA lattices, we projected the center of MA trimers radially onto the CA lattice layer and plotted the distribution of intersection points with reference to the center of the immature CA hexamer (Fig. 3i-l). Interestingly, while the overall distribution is relatively dispersed, there is a slight preference for the MA trimer to be in register with the CA trimer (or 3-fold axis) for WT VLPs (Fig. 3i,k). But more intriguingly, in L20K/E73K/A82T mutant VLPs, the intersection points became concentrated at the 3-fold axis, and the registration between the MA trimer and the CA trimer is much stronger than WT (Fig. 3j,l). This is consistent with the observation of a more ordered MA C-terminus in the L20K/E73K/A82T mutant MA compared to the WT (Fig. 3d). The L20K/E73K/A82T mutations indeed induce changes in the spatial organization of Gag, which may be attributed to the observed increased Gag processing in the L20K/E73K/A82T mutant (Fig 1a).

### MD simulations of immature MA trimer-lipid interactions

To understand the effect of L20K/E73K/A82T mutations on MA and lipid interactions, molecular dynamics (MD) simulations were employed to determine the regulatory effects of immature WT and L20K/E73K/A82T MA mutant proteins on lipid dynamics within the HIV-1 lipid membrane. The MD model encompassed an immature MA lattice (derived from PDBID 7TBP [40] and 2LYB [14] as indicated in the Methods section) embedded into an asymmetric lipid membrane of native HIV-1 lipidome composition (Fig. 4a, Table S2). We measured the instantaneous lateral displacement of lipids in the intravirion leaflet over 1 μs trajectories (Table S3, Simulations 4-6) using a vector field approach [41], revealing the formation of static regions in the membrane due to the presence of the MA lattice (Fig. S4, Movie S1). Notably, the average displacement over time indicated that these static regions in the intravirion leaflet correlate with the position of the MA lattice, suggesting that the MA lattice aggregates lipids in the intravirion leaflet (Fig. 4b). To further investigate specific MA regions related to aggregation of lipid headgroups, we conducted a comparative analysis of lipid contact occupancies between WT and mutant MA over the full trajectory. In both WT and L20K/E73K/A82T mutant MA, lipids aggregate with high lipid occupancies near positively charged amino acids, particularly at the HBR due to interactions with R21, K25, K26 and K29, as well as near helix 2 via interactions with R38 and R42 (Fig. 4c-d). Further comparative analysis revealed significant differences in lipid contact occupancies, with the L20K/E73K mutations increasing average occupancy at binding sites on the HBR and helix 2 and helix 4 (Fig. 4e). This suggests that L20K/E73K MA mutations enhance lipid-protein contact occupancies, consistent with its increased positive electrostatic potential on the lipid binding surface (Fig. 3e-f).

**Figure 4.**
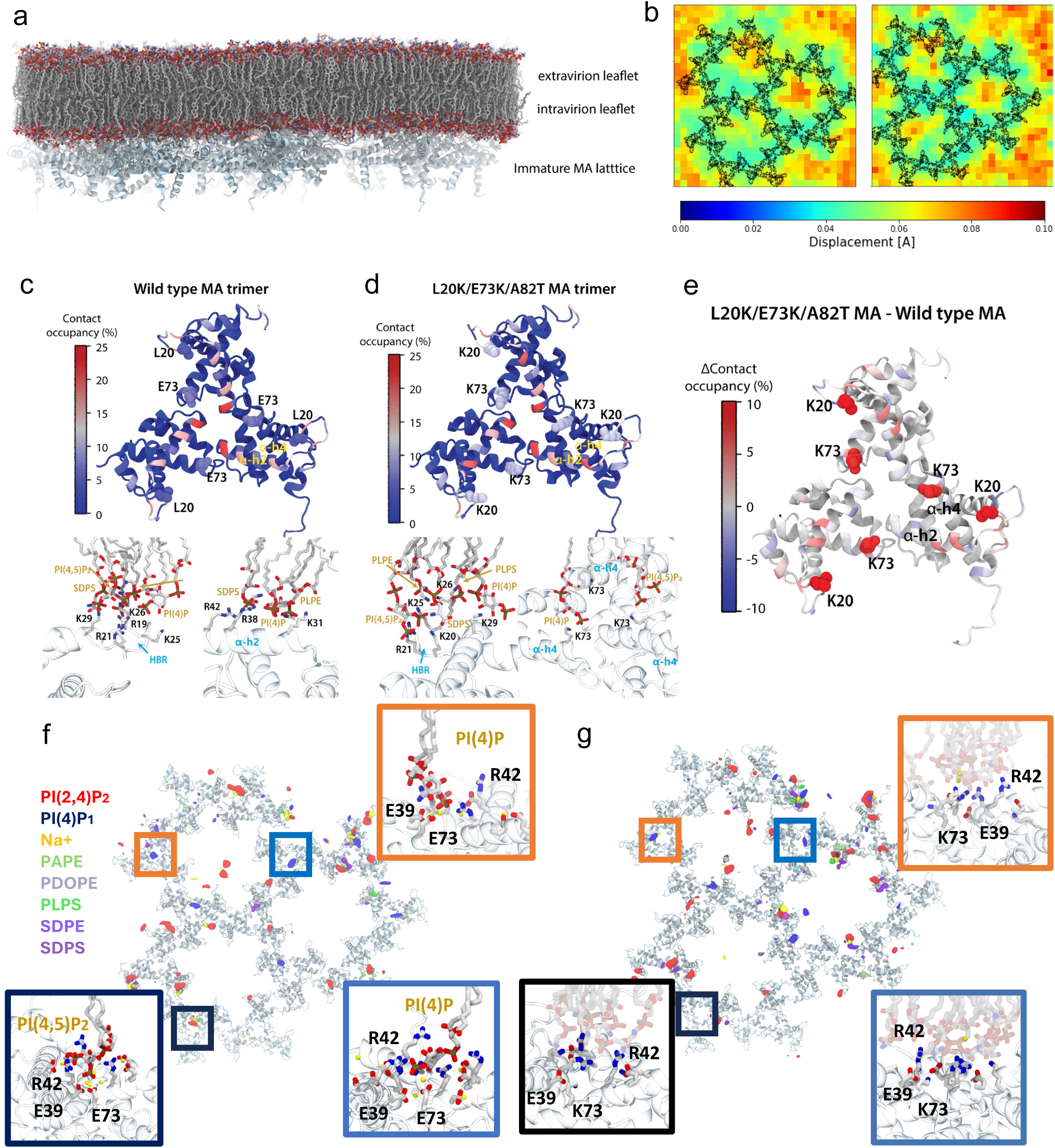
Lipid interactions with the immature MA lattice from molecular dynamic simulations. a) Model of the immature WT MA lattice embedded in an asymmetric lipid membrane of native HIV-1 lipidomics composition. b) Time average map of lipid displacement over 1 μs, in the presence of the immature WT MA lattice (left) or the immature L20K/E73K/A82T mutant MA lattice (right), average MA position over the trajectory is colored in black. c) (top) Wild-type MA trimer colored by lipid contact occupancy. (bottom) Protein-lipid interactions of positively charged regions in the HBR and helix 2 with intravirion leaflet lipids. d) (top) L20K/E73K/A82T MA trimer colored by lipid contact occupancy. (bottom) Interaction of residues at the HBR and E73K from helix 4 with intravirion leaflet lipids. e) L20K/E73K/A82T MA trimer colored by differences in lipid contact occupancy with respect to WT MA. f-g) Per-lipid occupancy maps for the intravirion leaflet lipid headgroups in the presence of WT MA, where E39 and E73 coordinate Na+ ions and R42 forms salt-bridges with PI(4,5)P_2_ and PI(4)P at trimer centers (f) and L20K/E73K/A82T MA where the protein-lipid interactions at trimer centers are only coordinated by R42 (g).

In addition, to elucidate the differences in the cryoET density between WT and mutant MA, we calculated the occupancy of lipid headgroups in the intravirion leaflet. From these calculations, we observed regions of high occupancy near trimer centers in the WT MA that were absent in the occupancy maps for mutant MA. Conversely, mutant MA displayed greater aggregation of lipid headgroups near the HBR and helix 4 (Fig. S5).

Further analysis of occupancy maps by lipid species indicated that the density at the trimer centers corresponds to specific binding of PI(4,5)P_2_ and PI(4)P lipid headgroups, while lipid aggregation in other regions appeared nonspecific (Fig. 4f-g). In fact, MD simulation trajectories reveal that at the trimer centers, E39 and E73 aggregate and coordinate Na^+^ ions with the phosphate groups of the PI(4,5)P_2_ and PI(4)P headgroups. This ion coordination, along with salt-bridge interactions with R42 stabilize PIPs at the trimer centers of the WT MA lattice (Fig. 4f). In the L20K/E73K/A82T mutant MA, however, E73 is substituted with K73, which forms salt-bridges with E39, E41 and transiently with other lipid headgroups but does not contribute to Na^+^ ion coordination. As a result, interactions at the trimer centers in mutant MA are only mediated by salt bridges with R42, making them less stable compared to WT MA (Fig. 4g, Movie S2).

### The L20K/E73K/A82T mutant does not form a mature MA lattice

As both CA and MA undergo major structural rearrangements upon PR-mediated Gag cleavage, converting from the immature to the mature configuration [18, 42], we next investigated the effect of L20K/E73K/A82T mutations on the structural maturation of VLPs. Using the same strategy employed to solve the structure of the MA layer in immature particles (Fig. S1), we determined the structure of the mature WT MA trimer at 7.3 Å resolution by subtomogram averaging (Fig. S6a-d, Table S1). The resulting structure overlaps closely with the previously published mature MA trimer structure, as well as the mature MA trimer lattice (Fig. S6e-f) [18]. In contrast, the mature L20K/E73K/A82T MA did not yield consistent subvolumes either through template matching or template-free approaches. Considering that no apparent MA density was observed at the IL of the viral membrane in raw tomograms of mutant VLPs (Fig. 2i-k), we attempted to locate the MA density layer using a density profiling approach (Fig. S7). By analyzing more than 20 complete VLPs, the radial density distribution profiles were aligned to the density valley of the outer leaflet (OL) and plotted along the radial distance with standard deviations (Fig. S7e). The distance of each density valley from the OL could then be calculated (Fig. S7d-f). To validate this method, we carried out such measurements using immature VLPs from both WT and the L20K/E73K/A82T mutant. Consistent with the tomography analysis, the radial density distribution profile from the immature mutant VLPs matches well to the profile from immature WT VLPs (Fig. 5a-c). There were no significant differences for the radial positions of MA, NC, and very small changes (∼ 3 Å) for N-terminal domain (NTD) of CA, C-terminal domain (CTD) of CA between the WT and the mutant (Fig. 5d). Interestingly, while mature WT VLPs showed the same MA localization as immature WT VLPs (Fig. 5e-g, IL/MA position), the mature L20K/E73K/A82T mutant VLPs displayed two separate density valleys. The first density valley corresponds to the IL of the membrane, which is located slightly closer (∼7 Å) to the outer leaflet compared to the IL/MA layer in WT (Fig. 5g-h). The second density valley, located ∼77 Å (STD: 4.7 Å) from the outer leaflet, was not present in mature WT VLPs (Fig. 5g-h, MA). These results suggest that the L20K/E73K/A82T mutant MA may have detached from the viral membrane upon maturation.

**Figure 5.**
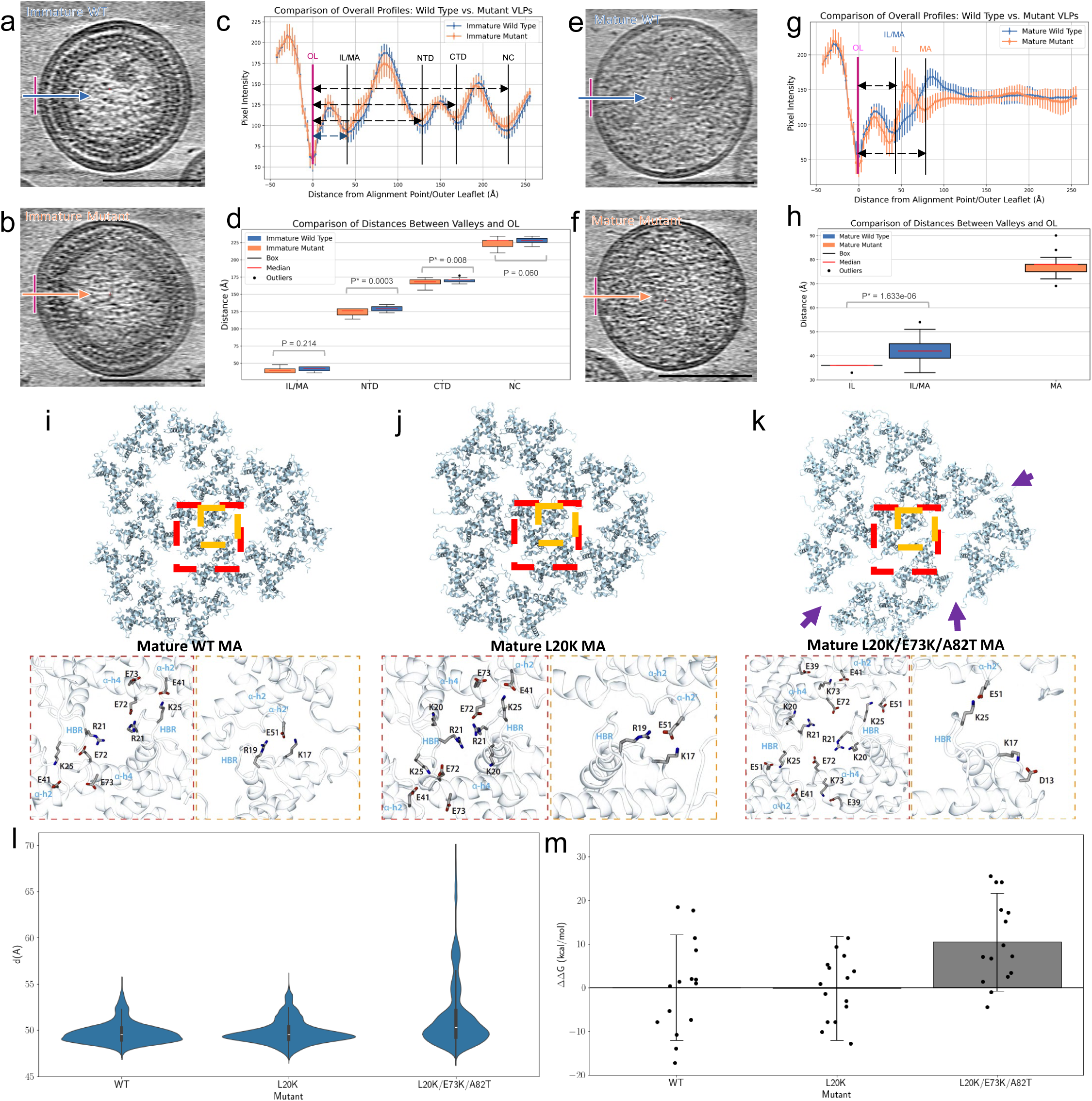
Analysis of mature MA lattice. a-d) Radial density profiles and distances of immature WT (blue) and mutant (orange) VLPs, measured from the outer leaflet (OL) of viral membrane. Profiles are aligned to the OL (the first valley). Representative central slices of immature WT (a) and L20K/E73K/A82T mutant (b) VLPs, enhanced by summing 10 neighboring slices, are demonstrated. Comparison of radial density profiles and distances between valleys and OL are plotted (c-d), revealing similar profiles and stable Gag protein organization during membrane assembly. Slight differences in the distances from IL/MA, NTD, CTD and NC to OL suggest that the mutant Gag shifts slightly towards IL. This indicates a potential increase in membrane binding affinity for the mutant. e-h) Radial density profiles and distances of mature WT and L20K/E73K/A82T mutant VLPs, measured from the outer leaflet of viral membrane. Representative central slice of mature WT (e) and mutant (f) VLPs, enhanced by summing 10 adjacent slices, are presented. Comparison of radial density profiles and distances between valleys and OL are plotted (g-h), displaying an additional valley further away from the outer leaflet in mutant VLPs. Each analysis includes over 20 complete VLPs per sample, with defocus values ranging from 4 to 5 um. Scale bar 100 nm. i-k) Mature MA lattice and inter-trimer interactions for WT MA (i), L20K MA (j) and L20K/E73K/A82T MA (k). l) MA trimer-trimer distance distributions through lattice perturbation MD simulations. m) Mature MA intertrimer estimated binding affinity compared to WT MA trimer interfaces via MM-GBSA calculations.

These findings prompted us to examine virions produced by the original single MA mutant L20K. The L20K mutation results in impaired endogenous reverse transcriptase activity and reduced viral infectivity [34, 36]. We produced and imaged immature and mature L20K mutant VLPs by cryoET as described above for the WT and the L20K/E73K/A82T revertant (Table S4). Interestingly, both immature and mature L20K VLPs exhibit a clear MA lattice at the IL of the membrane (Fig. S8a-f, brown arrowheads). Therefore, the loss of MA lattice appears to be a feature of the mature L20K/E73K/A82T mutant.

### MD simulations of the mature MA trimer lattice

To explore the effect of L20K/E73K/A82T mutations on the mature lattice, we turned to MD simulation. First, an atomistic model for an extended MA lattice was derived for WT MA, L20K MA, and L20K/E73K/A82T MA from the cryoEM/ET structures presented herein as described in the supplemental material. To simulate the stability of the mature WT MA and mature mutant MA lattices, we quantified the structural defects in the MA lattice through perturbation MD simulations in which we progressively adsorbed the mature MA lattice into the membrane, effectively varying the chemical environment (Table S3 Simulations 7-9; Movie S3). During simulations, we measured the trimer-trimer distances for every pair of neighboring MA trimers (Fig. S9) and quantified lattice structural stability by analyzing the distribution of trimer-trimer distances for MA trimers at the outer edge of the lattice. The latter is due to inner trimers being restrained by their neighbors.

Analyzing the trajectories resulting from our MD simulations of the MA lattice structure during the perturbative simulations (Table S3, Simulations 7-9), we observed notable alterations at the trimer-trimer interface in mature lattices due to the L20K/E73K/A82T mutations (Fig. 5i-k). Specifically, the mature WT MA lattice is typically stabilized by a network of inter-trimer salt bridge interactions involving residues from the HBR and glutamates from helices 4 and 2, namely K17-E51, R19-E51, R21-E72, K25-E41 (Fig. 5i). These salt bridges contribute significantly to the structural integrity and stability of the lattice. Although the L20K MA lattice maintains the same inter-trimer interactions as WT MA (Fig. 5j), the L20K/E73K mutations disrupt these critical inter-trimer salt bridges while promoting intra-trimer interactions, notably K73-E41, K73-E39, R21-E72, and K17-D13, which significantly alter the network of stabilizing forces within the lattice. Additionally, these mutations result in the displacement of the K25-E41 interaction to a new configuration between K25-E51 (Fig. 5k. Fig S10, Movie S4).

Comparing the trimer-trimer distance distributions aggregated over six replicates of the perturbation/adsorption simulations, we observe that the distributions for WT MA and L20K MA lattices are similar according to the Jensen-Shannon distance of 0.07 [43, 44]; the Jensen-Shannon distance evaluates the similarity between probability densities with a value between 0 and 1, 0 being identical and 1 most dissimilar. In contrast, the L20K/E73K/A82T MA lattice exhibited breaking events, with inter-trimer distances extending beyond 60 Å. This results in a multi-modal distribution that differs significantly from that observed for WT and L20K MA lattices, with a Jensen-Shannon distance of 0.30 (Fig. 5l). Additionally, tracking the occurrence of breaking events through the perturbation simulations revealed that the L20K/E73K/A82T MA presented a broken lattice on average 60% of the time compared to 17% of the time for WT MA and 6% for L20K MA (Fig. S9d, Movie S5). These results suggest that the WT and L20K mutant form more resilient MA lattices when compared to the L20K/E73K/A82T mutant, consistent with the result from cryoET.

Furthermore, we estimated the energetic favorability of the mutations for the self-association of MA trimers by calculating the binding affinity of trimer pairs throughout the simulations using the molecular mechanics generalized born surface area MM-GBSA method [45–47]. While L20K MA shows a negligible difference in binding affinity compared to WT MA trimer interface (ΔΔ*G* = 0.2 kcal/mol), the L20K/E73K/A82T mutations result in a more energetically unfavorable inter-trimer interface, with an average energetic penalty of Δ*G_L20K/E73K/A82T_* -Δ*G_WT_ =* 10.4 kcal/mol (Fig. 5m). As a result, we conclude that the L20K/E73K/A82T MA mutant is less likely to form higher-order trimeric assemblies.

These alterations not only compromise the stability of the mature MA lattice but may also explain why the mature L20K/E73K/A82T mutant MA appears to detach from the viral membrane. The enhanced intra-trimer interactions in the mutant further provide an explanation for the dimer and trimer observed for the MA mutant by SDS-PAGE (Fig. 1a-c). Interestingly, the purified WT and L20K/E73K/A82T mutant MA proteins display distinct features. Not only does the purified L20K/E73K/A82T mutant MA exist in monomer, dimer and trimer forms in the SDS-PAGE (Fig. S11a-c), it also forms filamentous structures seen in cryoEM micrographs, in contrast to the WT MA (Fig. S11d-e).

### Mutational analyses of the mature MA trimer-trimer interface

Our previous studies demonstrated that MA mutations, such as MA-R19L, do not significantly impair viral replication in T-cell lines or monocyte-derived macrophages despite being present at the putative mature MA inter-trimer interface [48]. This finding supports the results reported above for the L20K/E73K/A82T mutant that disruption of the mature MA lattice may not affect virus replication or infectivity. To further explore this question, we introduced alanine mutations at R19, E41, and E51 to disrupt the mature MA trimer-trimer interaction. Single-cycle infectivity and multi-cycle replication kinetics analysis demonstrated that these alanine mutations do not impair infectivity or viral replication (Fig. 6a-b).

**Figure 6.**
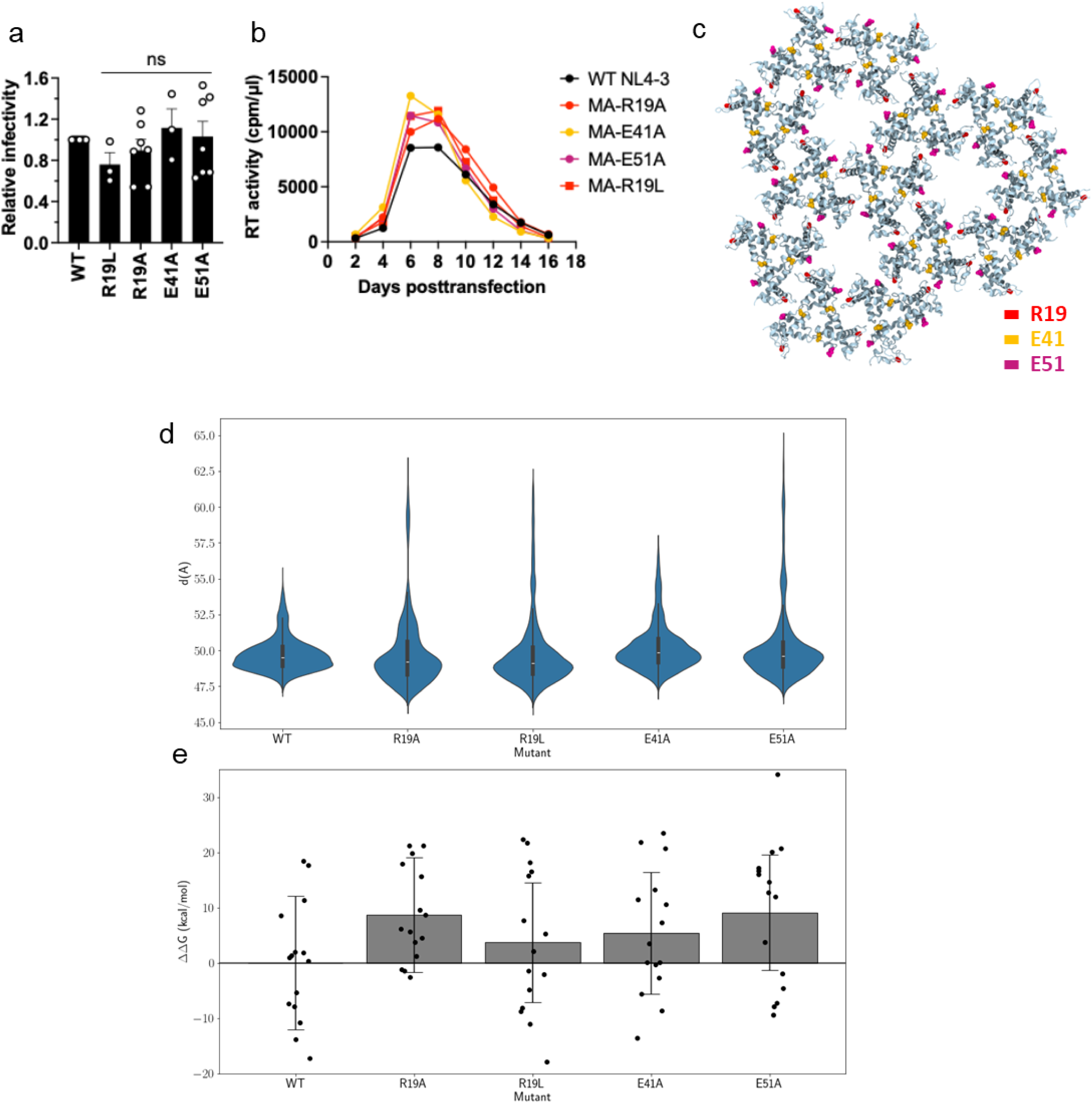
Mutations at the inter-MA trimer interface do not impair HIV-1 infectivity. a) Single-cycle, cell-free viral infectivity of the indicated MA mutants. RT-normalized virus stocks were used to infect TZM-bl cells. Luciferase activity was measured at 48 h post-infection. Relative infectivity is shown, normalized to 1 for WT NL4-3. Data from at least three independent experiments are shown as means ± SEM. ns, not significant, one-sample t-test. b) Replication kinetics of the indicated MA mutants. The H9 T-cell line was transfected with WT NL4-3 or the indicated MA-mutant proviral clones. Virus replication kinetics were monitored by measuring RT activity at the indicated time points. Data are representative of three independent experiments. c) Mature MA lattice showcasing the position of the mutations in the MA inter-trimer interfaces. d) MA trimer-trimer distance distributions for each salt-bridge-disrupting mutation R19A, R19L, E41A and E51A through 6 replicas of MA lattice perturbation MD simulations. e) Estimated mature MA intertrimer binding affinity compared to WT MA trimer interfaces via MM-GBSA calculations.

To assess the effects of these mutations on the mature MA lattice, we performed MD perturbation simulations on the mature MA lattices containing the R19A, R19L, E41A, and E51A mutations (Fig. 6c, Table S3, Simulations 10-13). We observe that the trimer-trimer distance distributions for all these mutants extend beyond the expected range for WT MA, differing from the WT trimer-trimer distance distribution with Jensen-Shannon distances of 0.33, 0.32, 0.18 and 0.24 for the R19A, R19L, E41A, and E51A mutations, respectively. This suggests that the MA trimers separate from one another in lattice-breaking events (Fig. 6d). Moreover, the simulations suggest that the trimer-trimer interfaces in these mutant MA lattices also become energetically unfavorable compared to those of the WT MA lattice, with the average binding affinity incurring penalties of 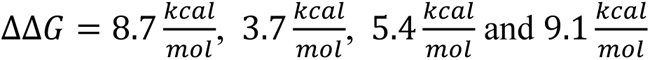 for R19A, R19L, E41A, and E51A, respectively (Fig. 6e). Although the effects of these mutations on the MA lattice are not as pronounced as those seen in the L20K/E73K/A82T mutant, the salt-bridge-disrupting mutations are predicted to compromise the MA lattice both structurally and energetically.

## Discussion

The HIV-1 MA domain plays a critical role in Gag targeting to the plasma membrane and Env incorporation into virions. Structural studies have shown that MA trimers form a hexagonal lattice on virions, with its organization influenced by PR-mediated Gag cleavage [18]. However, the biological significance of this structural rearrangement in the MA lattice following Gag cleavage remains unclear. In this study, we compared the structure of the immature and mature MA lattices in WT and L20K/E73K/A82T MA mutant virions using biochemical, structural, and computational approaches. Our results demonstrate that while the L20K/E73K/A82T MA mutant exhibits distinct biochemical MA properties compared to WT, it maintains similar levels of infectivity in the H9 T-cell line and in PBMCs [32]. Structural and computational analyses further reveal that the L20K/E73K/A82T mutant lacks a well-organized mature MA lattice in virions due to unstable inter-trimer interactions despite forming a WT-like immature MA lattice. Additionally, mutations in MA residues predicted to contribute to inter-trimer interactions in the mature MA lattice did not affect viral infectivity or viral replication. These findings suggest that a well-organized mature MA lattice is not essential for viral infection.

The HBR, which includes residues Arg21, Lys25, Lys26, Lys29, and Lys31, creates a positively charged area on the MA globular domain’s surface that facilitates membrane binding via electrostatic interactions with the acidic headgroups of phospholipids including PI(4,5)P_2_ and PS [26, 49, 50]. The L20K mutation introduces an additional positive charge within the HBR, thereby enhancing its membrane affinity [34]. NMR studies have demonstrated that when MA binds to phosphatidylserine (PS), there is a favorable interaction between the acidic headgroup of PS and the R42 residue of MA [14]. The crystal structure of the MA trimer reveals a central pore surrounded by three R42 residues [15]. The density observed in the trimer center of both immature and mature WT MA lattices indicates the possible presence of a negatively charged entity that could interfere with phospholipid interaction. In the immature L20K/E73K/A82T mutant, the absence of central density, coupled with upward reorientation of R42 in the AlphaFold2 prediction, as well as the E73K mutation, are presumed to enhance interactions with lipid headgroups, consistent with the role of R42 in phospholipid binding. The collective structural modifications induced by L20K, E73K, and the repositioning of R42 create a highly positive surface for phospholipid binding and docking, offering an explanation for the increased Gag binding affinity observed for the L20K/E73K/A82T mutant [32].

Recent structural analysis suggests that proteolytic cleavage of SP2 from the Gag precursor by PR is required to form a mature MA lattice [51]. The SP2 peptide was shown to bind to the inter-trimer interface, previously predicted to be a PI(4,5)P_2_ binding pocket [18], and contributes to stabilizing the mature MA lattice. This study also reported that fusion kinetics are influenced by the release of SP2. However, previous studies have shown that deletion of SP2 or modifications at the NC-SP2 cleavage site confer only minor defects in virus infectivity [52–54]. The results reported herein also show that formation of a mature MA lattice, presumably required for SP2 binding, is not strictly required for HIV-1 particle infectivity. Thus, while SP2 may contribute to the reorganization of the immature MA lattice to the mature form, further work will be required to fully elucidate the interplay between formation of the mature MA lattice, SP2 binding, and particle infectivity.

## Materials and Methods

### Cell lines and plasmids

HeLa cells (obtained from the American Type Culture Collection [ATCC]), human embryonic kidney (HEK) 293T cells (obtained from the ATCC), and TZM-bl cells (obtained from J.C. Kappes, X. Wu, and Tranzyme, Inc. through the National Institutes of Health [NIH] AIDS Reagent Program [ARP], Germantown, MD) were maintained in Dulbecco’s modified Eagle’s medium supplemented with 10% fetal bovine serum at 37°C in 5% CO2. The H9 T-cell line was cultured in RPMI-1640 medium supplemented with 10% FBS at 37°C in 5% CO_2_.

The full-length HIV-1 molecular clone pNL4-3 [55] was used in this study. pNL4-3 MA-L20K and L20K/E73K/A82T, and R19L were described previously [32, 34, 48]. We introduced alanine mutations in MA by HiFi DNA Assembly Master Mix (NEB) according to manufacturer’s instructions. For biosafety reasons, VLPs for structural analysis were prepared with a pNL4-3 derivative containing inactivating mutations in integrase (IN) and reverse transcriptase (RT) (pNL4-3 IN-D116N/RT-D186N) [56] or subviral plasmid pCMVNLGagPolRRE, which expresses NL4-3 GagPol from a CMV promoter [57]. To prepare immature VLPs, the PR active site mutation PR-D25N was introduced into pNL4-3 RT-D186N/IN-D116N or pCMVNLGagPolRRE plasmids. The MA-coding regions of pNL4-3 MA mutants were cloned into pCMVNLGagPolRRE using BssHII and SpeI restriction sites. The HIV-1 Env expression vectors, pIIINL4env and pIIINL(AD8)env were described previously [48, 58].

### Antibodies

Anti-HIV-1 Pr55Gag + p24 + p17 antibody (ab63917) and anti-tubulin antibodies (B-5-1-2) were purchased from Abcam and Sigma-Aldrich, respectively. Anti-gp41 (Chessie 8) and polyclonal anti-p17 antibodies (ARP-4811) were obtained through the National Institutes of Health (NIH) HIV Reagent Program. Additionally, a polyclonal anti-Gag p17 antibody (PAB1178) was obtained from Abnova.

### Preparation of viruses or VLPs

For VLPs used in the structural analysis, HEK293T cells were co-transfected with pCMVGagPolRRE, pIIINL4env, and pSV-Tat [59] at a ratio of 16:4:1 using GenJet In Vitro DNA Transfection Reagent (Ver. II) (SignaGen Laboratories). Alternatively, pNL4-3 RT-D186N/IN-D116N and pIIINL(AD8)env were co-transfected at a ratio of 10:1. Culture media from transfected cells were harvested at 48 h post-transfection and passed through a 0.45 µm polyvinylidene fluoride (PVDF) filter. The VLPs were concentrated by ultracentrifugation through an 8% OptiPrep density gradient (Sigma-Aldrich) (100,000 x *g*, Sorvall AH-629 rotor) for 1 h at 4 °C. The concentrated VLPs were further purified by ultracentrifugation (120,000 x *g*, Sorvall TH-660 rotor) through a 10 – 30% OptiPrep gradient for 2.5 h at 4°C. The opalescent band was harvested, diluted with PBS, and ultracentrifuged at 110,000× *g* at 4 °C for 2 h.

Pelleted particles were resuspended in 5% sucrose/PBS solution and stored at -80°C until use. For cryoEM grid preparation, a 3 µl aliquot was placed onto a glow-discharged lacy grid (AGS166-3 Lacey Carbon Films on 300 Mesh Copper Grids) and plunge-frozen in liquid ethane using the Leica EM GP 2 system for back-side blotting.

### Membrane stripping assay

The HEK293T or HeLa cell lines were transfected with WT or mutant pNL4-3 molecular clones using Lipofectamine 2000 (Invitrogen). Viruses were incubated with various concentrations of NP-40 for 5 min. Following incubation, the viruses were purified by ultracentrifugation through 20% sucrose cushions (60,000 x *g*) for 45 min at 4°C. For sedimentation analysis, the concentrated viruses incubated with 0.075% NP-40 were subjected to ultracentrifugation through a 10-70% sucrose gradient (60,000 x g) for 16 h. Alternatively, the viral membrane was stripped by a ‘spin-through’ detergent-treatment method with minor modification [37]. Briefly, the concentrated viruses were subjected to ultracentrifugation (60,000 × *g* for 2h at 4°C) through a layer of 1% Triton X-100 into a linear OptiPrep density gradient (10 – 60%). Aliquots from each fraction were lysed and subjected to western blot analysis.

### Western blotting

Cell-and virus-associated proteins were solubilized in lysis buffer (30 mM NaCl, 50 mM Tris-HCl pH 7.5, 0.5% Triton X-100, 10 mM iodoacetamide, complete protease inhibitor [Roche]). Lysates boiled in 6X loading buffer (7 ml 0.5 M Tris-HCl/0.4% SDS, 3.8 g glycerol, 1 g SDS, 0.93 g DTT, 1.2 mg bromophenol blue) were subjected to SDS–PAGE and transferred to polyvinylidene disulfide membranes (Merck Millipore). After blocking the membranes with Azure Fluorescent Blot Blocking Buffer (Azure Biosystems), the membranes were probed with the indicated antibodies (1:5000 – 10,000) at 4°C overnight and then incubated for 1 h with species-specific AzureSpectra Fluorescent Secondary Antibodies (Azure Biosystems). After the final washes, bands were detected by fluorescence with a Sapphire Biomolecular imager (Azure Biosystems). Quantification was performed using ImageJ software.

For western blotting of WT MA or MA mutant proteins (Supplemental Figure 11a-b), these proteins were first expressed in *Escherichia coli* (*E. coli*) Rosetta 2 (DE3) cells by inducing with 0.1 mM isopropyl β-D-1-thiogalactopyranoside (IPTG) at 37 °C for 3 h in Luria-Bertani medium. Following expression, cells were harvested by centrifugation and resuspended in lysis buffer (50 mM Tris-HCl, pH 8.0, 150 mM NaCl, 0.1% Triton X-100). The lysate was clarified by centrifugation at 20,000 x g for 30 minutes using an SW-32 rotor. Both the soluble supernatant and insoluble pellet fractions were analyzed via SDS-PAGE to assess the solubility of the expressed protein. Proteins were then transferred to a nitrocellulose membrane, which was probed with a polyclonal anti-Gag p17 antibody (PAB1178). After primary antibody incubation, the membrane was washed thoroughly and incubated with an anti-mouse horseradish peroxidase (HRP)-conjugated secondary antibody (Sigma-Aldrich, A0168) at a 1:5000 dilution for 1 h at room temperature. Protein bands were visualized using Clarity Western ECL substrate (Bio-Rad) according to the manufacturer’s protocol.

### Single-round infectivity assay

Single-round infectivity assays were performed as previously described [60]. The amount of virus in the supernatant was quantified by an RT assay, performed as described previously [60]. TZM-bl cells (1.0 × 10^4^ cells) in 96-well plates were incubated with RT-normalized virus stocks. At 48 h post-infection, luciferase activity was measured using the Britelite plus reporter gene assay system (PerkinElmer) and GloMax Navigator microplate luminometer (Promega).

### Virus replication kinetics analysis

Virus replication was monitored in H9 T-cell line as previously described with minor modifications [61]. H9 cells were incubated with the indicated pNL4-3 MA variants (1.0 µg DNA/1.0 × 10^6^ cells) in the presence of 700 µg/ml DEAE-dextran at 37°C for 15 min. Transfected cells were cultured in 24 well plates. Aliquots of supernatants were collected to measure RT activity, and cells were split 1:1 every other day with fresh medium.

### Protein purification and cryoEM

Wild type MA and MA L20K/E73K/A82T mutant genes were cloned into the pSY5 vector [62], which includes an N-terminal 8-histidine tag followed by a human rhinovirus (HRV) 3C protease cleavage site (LeuGluValLeuPheGln↓GlyPro). The plasmid was transformed into E. coli strain BL21 (DE3). The proteins were expressed overnight (18°C) in terrific broth medium supplemented with 0.4% (vol/vol) glycerol, following induction by 0.2 mM IPTG (isopropyl-D-1-thiogalactopyranodside) at a cell density characterized by an optical density at 600 nm (OD600) of 0.8–0.9. The resultant cell pellets (10 g) were resuspended in binding buffer 50 ml (20 mM HEPES, 500 mM NaCl, 20 mM imidazole and 1 mM TCEP, pH 7.7) supplemented with Triton X-100 (0.01% [vol/vol]), protease inhibitor cocktail (Set III, EDTA-free, Calbiochem) and benzonase (2 μl of 10,000 U/μl, Merck). Cell lysis was performed using an ultrasonic cell disrupter Vibra-Cell (Sonics). The proteins bound to a Ni-NTA affinity chromatography column (HisTrap FF GE Healthcare), washed with binding buffer containing 20 mM Tris–HCl (pH 8.0), 1 M NaCl, 1 M (NH4)2SO4, 30 mM imidazole, and 5 mM β-mercaptoethanol. The target protein was eluted with 15 mL of elution buffer composed of 20 mM Tris–HCl (pH 8.0), 500 mM NaCl, 1 M (NH4)2SO4, 5 mM β-mercaptoethanol, and 500 mM imidazole. The eluted fraction was concentrated to approximately 2 mL, and the His tag was cleaved with 0.1 mg/mL of His-tagged HRV 3C protease during an overnight dialysis against a buffer containing 20 mM Tris–HCl (pH 8.0), 500 mM NaCl, 5% (vol/vol) glycerol, and 1 mM dithiothreitol (DTT). Following cleavage, the reaction mixture was diluted to a total volume of 50 mL. The untagged MA protein, along with the His-tagged HRV 3C protease and uncleaved MA, were separated using reverse Ni-NTA affinity chromatography. The untagged MA protein was collected as the flow-through fraction. To denature the protein, 2 mL of the flow-through was combined with 10 mL of denaturation buffer (20 mM Tris–HCl, pH 8.0, 1 M NaCl, 1 M (NH_4_)_2_SO_4_, 1 mM DTT, and 6 M urea) and allowed to incubate overnight during dialysis. Following this, the sample was concentrated to a final volume of 1 mL, and size-exclusion chromatography (SEC) was performed using a Superdex 200 10/300 increase column (GE Healthcare, Little Chalfont, United Kingdom). The column was equilibrated with SEC buffer (20 mM Tris–HCl, pH 8.0, 150 mM NaCl, and 1 mM DTT) and run on an Akta FPLC system (GE Healthcare) at a flow rate of 0.2 mL/min. Peak fractions containing the refolded MA protein were pooled and concentrated (2000 MWCO Amicon® Ultra Centrifugal Filter concentrator) for cryoEM analysis.

Three microliters of purified WT or mutant MA protein (1 mg/mL) was applied to glow-discharged Lacey carbon-coated copper grids (300 mesh, Agar Scientific). The grids were blotted under 100% humidity and plunge-frozen into liquid ethane using a Vitrobot Mark IV (Thermo Fisher Scientific). Following vitrification, the grids were clipped and screened using a Glacios cryo-electron microscope (Thermo Fisher Scientific).

### CryoET and subtomogram averaging

The cryoET data were acquired with 300 kV Titan Krios (eBIC Krios III), utilizing Thermo Fisher Scientific Falcon 4i detector & Selectris X energy filter with 5eV window. Tomographic tilt series were collected with tilt angles ranging from -60° to +60°, in 3° increments, following a dose-symmetric scheme with a group size of 3, using Tomo 5 (Thermo Scientific). The effective magnification was set to 81,000x, resulting in 1.50 Å/pixel on the specimen. The nominal defocus for each tilt series ranged from 1.5 to 5.5μm. Movies were captured for each tilt, consisting of 10 frames in counting mode. Dose requirement is 3 e/Å² per tilt, resulting in total cumulative 123 e/Å² per series. Details of all data acquisition settings can be found in Table S1. Workflow for data processing is presented in Fig. S1. Frames were motion-corrected using MotionCor2 [63]. Fiducial alignment of tilt stacks was done using IMOD [64]. Subtomogram alignment and averaging were conducted using emClarity [39, 65] and Relion [38].

For CA of immature mutant WT and mutant particles, we selected particles from 6x binned, non-contrast transfer function (CTF)-corrected tomograms template matching through emClarity/1.5.0.2, employing a 28 Å low-pass filter on the EMD-8403 [66]. The magpiEM tool (https://github.com/fnight128/MagpiEM) was used for cleaning based on the geometric restraints of the CA lattice. 3D alignment and averaging for hexamer CA was incrementally refined from 6x to 1x binning in emClarity/1.5.3.10, preserving C6 symmetry.

For MA of immature mutant WT and mutant particles, the immature lattice structure from PDB 7OVQ [18] served as the initial template. We generated an extended 400Å-wide lattice map for MA template matching on 6x binned, CTF-corrected tomograms using emClarity/1.5.3.10, with a 28 Å low-pass filter applied. The magpiEM tool was also utilized for data cleaning based on geometric constraints, particularly between the bilipid and CA layers. Top view subtomograms, identified by a tilt angle under 45° relative to the beam, were selected for iterative alignment, beginning from 6x binning down to 4x binning. Subsequently, these subtomograms were subjected to global in-plane and local out-of-plane refinements at their 4x binned top view alignment. This refined positioning also included further cleaning to remove duplicates and ensure adherence to local MA lattice constraints. The further cleaned MA subtomograms were then prepared for averaging in Relion 4, enforcing C3 symmetry after global refinement at bin4 and local refinement at bin2.

For mature WT MA, we utilized a mature, 400Å-wide lattice model extended from PDB 7OVR [18] as a template for matching on 6x binned, non-CTF-corrected tomograms using emClarity/1.5.0.2 with a 28 Å low-pass filter to the template. The magpiEM script was used to select MA subtomogram matches based on geometric constraints beneath the bilipid layer. Criteria for further selecting trimeric MA structures ensured the exclusion of unreliable peaks. This selection step prepared the dataset for further refinement in Relion 4.0.0, where 3D refinement was performed at gradually decreasing bin levels, maintaining C3 symmetry throughout.

### MA lattice arrangement analysis

In the comparative analysis for subtomograms that were mapped back, we define the distance and tilt angle for trimer-trimer pairings within the MA lattice. The lattice maps of CA hexamers and MA trimers were generated using the ArtiaX for UCSF Chimera X [67, 68]. To facilitate a reasonable comparison between the immature mutant and WT MA lattice maps, the density map threshold for MA is calculated using a 1% false discovery rate [69].

### Radial density profile analysis

We enhanced the signal from the central slice of a VLP by summing it with 10 neighboring slices. The positions of the OL within the slice were identified to serve as a reference point for subsequent analysis. To accurately locate the OL, we manually marked points along its perimeter, fitted these points to a circle path, and generated normal line density profiles perpendicular to this path. The position of the OL in each profile was determined by identifying local minima around the plotted circle. These profiles were then averaged, aligning them based on the OL position to ensure consistency. We accumulated a substantial number of VLP average profiles (over 20 VLPs for each sample) to ensure the robustness of our analysis. The final average profile was displayed with error bars to illustrate the membrane-associated density representation of the sample. Additionally, a box plot of the distance between different valleys and the OL was created to further analyze the data (Fig. S5).

### HIV-1 lipid membrane modeling

In preparation for the atomistic molecular dynamics simulations of an MA lattice embedded in an HIV-1 lipid membrane with native composition, we modeled the model membrane by the following procedure: First, we derived the stoichiometric ratios of lipid species corresponding to native HIV-1 lipidomics [70]; for the purpose of modeling an asymmetric membrane, the intravirion and outervirion leaflet ratios for each lipid species were estimated from united atom-based simulations of an HIV-1 lipid vesicle [71]. The resulting stoichiometric ratios between different lipid species are shown in Supplementary Table S2.

We previously showed that the viral membrane of a spherical virion is asymmetric due to the differences in curvature between the outer and inner leaflets of the virion [71]. Therefore, to build a flat asymmetric lipid bilayer of native HIV-1 lipid composition we followed the procedure proposed by Pastor and co-workers [72]. For the latter, we prepared two symmetric lipid bilayers using CHARMM-GUI [73, 74]: a) a lipid bilayer corresponding to the extravirion leaflet composition (Table S2) and b) a lipid bilayer corresponding to the intravirion leaflet composition (Table S2). Each symmetric membrane was solvated and ionized by adding NaCl ions to neutralize the charge of the system and further solvated in a water box containing a salt concentration of 150 mM NaCl. The resulting symmetric membranes were then equilibrated following the CHARMM-GUI membrane builder protocol [75, 76]. Namely, the system was minimized using a conjugate gradient scheme until the gradient converged below 10 kcal/mol/Å^2^, followed by a lipid tail melting procedure where the lipids molecules were allowed to reorganize and repack the membrane for 2 ns at constant temperature of 310 K, constant pressure of 1 bar, and containing 905,725 and 831,950 atoms for the intravirion and extravirion symmetric leaflet systems, respectively; water molecules are kept out of the membrane hydrophobic core by applying external forces on the oxygen atoms of water molecules that penetrate in the membrane. Subsequentely, the system is equilibrated for 50ns at 310 K, 1 bar and containing over 830,000 particles; constant area pressure control was employed with a total cross-sectional area of 100,997 Å^2^ and 86,201 Å^2^ for the intravirion and extravirion symmetric leaflet systems, respectively; symmetric membrane simulations were performed in NAMD 2.15b14 (Table S3, Simulations 1-2). All membrane molecular dynamics simulations used the hydrogen mass repartition scheme [77], which enables a 4 fs timestep for time propagation. During equilibration simulations, temperature was maintained at 310 K using a Langevin thermostat with a thermal coupling constant of 1 ps^-1^ and a pressure of 1 atm via a Nose-Hoover barostat with a period of 100 fs and decay time of 50 fs.

Following equilibration of the symmetric lipid bilayers, the coordinates of the lipid molecules, ionic molecules, and water molecules associated with the inner leaflet of the intravirion symmetric membrane and the outer leaflet of the extravirion symmetric membrane were extracted and merged into a single asymmetric membrane (following the work of Pastor and co-workers [72]). The latter procedure conserves the ion and water molecules absorbed to lipid headgroups in each leaflet, as well as water solvation shells, in the asymmetric membrane. Due to the differences in lipid per area and lipid compositions in the intravirion and extravirion leaflets, the intravirion leaflet resulted in a larger area than the extravirion leaflet and was thus trimmed to fit the dimensions of extravirion leaflet while maintaining the molar fraction ratios for each lipid species. The resulting asymmetric membrane extended over an area of 29.4 nm x 29.4 nm with a resulting composition detailed on Supplementary Table S2.

After building the HIV-1 flat asymmetric lipid membrane, we performed lipid tail melting and equilibration following the aforementioned procedure implemented for the symmetric membranes [75]. Subsequently, we performed three independent replicates of 1 μs canonical MD simulation in the NPT ensemble at a temperature of 310 K and pressure of 1 atm using the Langevin thermostat and Nose-Hoover barostat with the same parameters as above, keeping the ratio of X-Y unit cell constant while allowing fluctuation in all axes. Long-range electrostatics were calculated using the particle mesh Ewald method [78] with a short-range cutoff of 12 Å and switching parameter of 10 Å (Table S3, Simulation 3). All simulations were performed using NAMD 2.15alpha2 and NAMD 3beta6 [79].

### MA lattice modeling

A model of the full-length MA was derived from the X-ray crystal structure of the MA trimer (PDBID 7TBP) [40], complemented with the terminal MA residues 109 to 115 from the NMR myrMA monomer structure (PDBID 2LYB) [14]. Thirteen MA trimers were rigid-body fitted into EM densities for the WT immature MA, L20K/E73K/A82T immature MA and WT mature MA densities to build the respective MA lattice models; each model consisting of a 12-mer of trimers (Figure 5i). The three systems were then prepared for molecular dynamics simulations by adding hydrogens to MA according to the protonation state of the amino acids at pH 7.0 as predicted by propKa3 [80, 81]. The protein was then solvated with TIP3P water molecules into a periodic box and ionized with Na^+^ and Cl^-^ ions to achieve a concentration of 150 mM and neutral charge using the *solvate*, *cionize* and *autoionize* plugins in the Visual Molecular Dynamics (VMD) software [82].

The rigid body fitted MA lattices were then refined via Molecular Dynamics Flexible Fitting (MDFF) [83] by running molecular dynamics in a constant number of particles, pressure and temperate (NPT) ensemble and utilizing the cryoEM density as a grid-based biasing potential coupled to the protein heavy atoms, effectively biasing their movement to fit the electron density. MDFF was performed for 10 ns with a coupling gird scaling factor of 0.3 a.u. L20K MA mature and immature lattices, as well as the initial model for the L20K/E73K/A82T MA mature lattice simulations, were built from the mature or immature WT MA lattices by introducing the mutations with the Mutator plugin in VMD [84]. The structure of each MDFF-fitted MA lattice was further refined utilizing an automated structure refinement procedure[85] using RosettaScripts[86] as preparation for deposition to the protein data bank. Final refined structures are deposited with accession codes PDB ID 9EK3 and PDB ID 9EK2 for the immature WT and L20K/E73K/A82T MA assembly, respectively, and PDB ID 9EK1 for the mature WT MA assembly.

### MA embedded in an authentic HIV-1 lipid membrane complex

After deriving models for MA lattices guided by the electron density, we utilized the protein coordinates and merged them with the equilibrated coordinates of the previously prepared HIV-1 lipid membrane. First, we placed the MA lattice 10 nm from the intravirion leaflet and outside of the simulation box; subsequently we solvated the MA lattice and ionized with Na^+^ and Cl^-^ ions to a 150 mM concentration. Then, to embed MA into the lipid membrane, we applied a constant velocity pulling to the last carbon of the MA myristoyl tail towards the hydrophobic core of the lipid bilayer at a rate of 5 Å/ns by applying successive harmonic restraints with a spring constant of 0.5 kcal/mol/Å^2^ in 100 successive 0.1 ns MD simulations until all myristoyl tails were embedded in the lipid bilayer hydrophobic core. These pull and wait simulations [87, 88] were performed in an NPT ensemble with constant area pressure control and a pressure of 1 atm via a Nose-Hoover barostat with a period of 200 fs and decay time of 100 fs, while the temperature was maintained at 310K using a Langevin thermostat using a high Langevin coupling constant of 100 ps^-1^ to dampen the dynamics of the lipid membrane during myristoyl tail insertion. This effectively restricted the axial movement of the lipid membrane without adding constraints to the positions of the lipid headgroups.

After the myristoyl tail insertion procedure described above, the Langevin coupling constant was reduced to 10 ps^-1^ over 20 ns, and subsequently to 5 ps^-1^ and 1ps^-1^ over the same period. This approach reduced drag and allowed the MA lattice to be adsorbed into the lipid membrane without restricting protein and lipid dynamics. This procedure was applied to all MA lattices to build mature and immature MA-membrane complexes for wild-type MA, L20K and L20K/E73K/A82T mutant MA, resulting in six MA-membrane complex models indicated in Supplementary Table S3 (Simulations 4-9) as well as the four salt-bridge perturbing MA mutations R19A, R19L, E41A and E51A (Simulations 10-13).

### Protein-lipid interaction analysis

Each immature MA-membrane complex was subjected to three replicates of canonical MD simulations in an NPT ensemble for 1 μs (Table S3 Simulations 4-6). Throughout each trajectory, we tracked the instantaneous lateral displacement of lipids in the intravirion leaflet using a vector velocity field [41]. This method discretizes the space into a grid of 2D voxels and tracks the movement of lipids though the membrane by calculating the displacement of the center of mass of the group of lipids 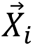 in the i-th voxel from a time *t* to a time *t + dt* as a vector

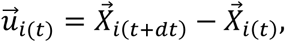

which is then visualized using streamlines. We used the lipid headgroup atoms to calculate centers of mass on each voxel a grid spacing of 10Å to which ensured an average of ∼5 lipid molecules per voxel and a *dt* = 0.2 *ns* as indicated in the work by Chavent, et.al. [41]. As indicated in aforementioned work, to obtain a smooth transition of velocity vectors through time, we preprocessed the 1 μs trajectories using a low-pass filter of the coordinates using the *gmx_filter* tool in GROMACS version 2024.3 [89]. Time averages of lipid displacement are then calculated on a per-voxel basis to identify static regions in the membrane.

Lipid-protein contacts were analyzed throughout the simulation in VMD [82] using in-house Tool Command Language [90] scripts. These contacts were measured over each frame of the simulation trajectory using a distance threshold of 3.5 Å between protein and lipid headgroup heavy atoms. Contact occupancies were measured for each MA residue and every lipid species in the intravirion leaflet of the membrane and averaged over the 39 monomers in the 12-mer of trimers assembly. This approach ensures that if a lipid-protein interaction has high occupancy in one monomer but is not present in the others, its average contribution to the MA-membrane interactions is reduced. The highest MA-lipid occupancy for each MA residue was then represented visually in the protein structure model of an MA trimer (Figure 4f-h), permitting the identification of high occupancy regions in both the WT MA lattice and L20K/E73K/A82T MA lattice.

Lipid headgroup occupancies were calculated for all lipid species and per-lipid species through the 1 μs trajectories (simulations 4-6) with a spatial resolution of 1Å by utilizing the volmap pluigin in VMD[82] and the voltool plugin was used to generate a difference map between the lipid headgroup occupancies in the L20K/E72K/A82T MA and WT MA simulations. Occupancy maps are visualized at the 0.25 isovalue representing sites where lipid headgroups localize for over 25% of the simulation.

### Perturbative MD simulations of MA lattices adsorption by lipid bilayers

The stability of mature MA lattices was probed in a novel perturbative approach based on controlled damping of Langevin dynamics during protein-membrane adsorption, as described below. The main idea of our method consists of perturbing the molecular environment surrounding MA, by transferring MA from a liquid solvent (water and ions) to a complex lipid environment. To control the temperature in MD simulations, the movement of particles in the system is coupled to a thermal bath at a constant temperature T by utilizing a thermostat. In the case of the widely-used Langevin thermostat [91], the dynamics of the *i*-th particle in the ensemble, with mass *m_i_* and position 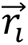 is described by

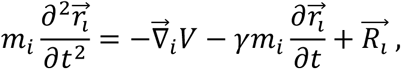

where *V* is the molecular potential, 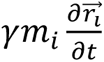 is a dissipative drag force with friction coefficient γ, and 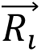 is a random force due to thermal motion of the particles in the coupled thermal bath, modelled as a gaussian noise with zero mean and variance *σ*^2^ = 2*m_i_γk*_B_T/Δt. The dissipative drag force dampens the momentum of particles and the friction coefficient γ effectively controls the viscosity of the solvent. If γ is too large, the system is overdamped and obeys Brownian dynamics. In contrast, if γ is too small, the system is underdamped and may have trouble dissipating heat in a short simulation time; with the limit case γ=0 representing a system with no temperature coupling.

The Langevin thermostat guarantees ergodicity and -in equilibrium simulations-evolution towards canonical ensemble distributions after enough sampling regardless of the friction coefficient [92]. However, previous computational studies have shown that in non-equilibrium processes, variations in the Langevin friction coefficient can cause differences in macroscopic observables (material hardness [93], diffusion coefficients [94]) and microscopic structure (crystal lattice plastic deformation [93]).

Protein-membrane adsorption is a dynamic process in which the both protein and lipids can undergo structural changes [95, 96]: localization of protein can induce the formation of lipid raft domains in the membrane [97, 98] while protein conformational changes can be promoted by adsorption into the membrane [99–101]. For HIV-1 MA, protein adsorption is driven by electrostatic interactions between negatively charged lipid headgroups and positively charged residues in the surface of MA and is enhanced by the hydrophobic interactions caused by myristoil-tail insertion in the membrane [102, 103].

In our approach, we conduct non-equilibrium MD simulations of MA adsorption to the lipid membrane by gradually decreasing the Langevin friction coefficient in three successive stages from 10 ps^-1^ to 5 ps^-1^ to 1 ps^-1^, each lasting 20 ns. This method maintains a constant temperature and ensure proper heat dissipation throughout the adsorption process while the sequential reduction in the friction coefficient allows for the gradual displacement and interaction of the MA lattice with the lipid membrane via electrostatic interactions. In this process, as the mature MA lattice approaches the membrane, it is gradually perturbed by the changes in chemical environment and decreasing drag forces until it reaches the adsorbed-equilibrium conformation. Structural stability of the MA lattice through the simulation depends on the strength of interactions between MA trimers and their ability to withstand the variations in chemical environment and drag forces.

Each mature MA lattice was probed via 6 replicas of perturbation/adsorption experiments (Table S3 Simulations 7-13) in which the L20K, L20K/E73K/A82T, R19A, R19L, E41A or E51A mutations were introduced to the mature WT MA system after embedding the myristoyl tail in the membrane and reionizing the system to charge neutrality. The membrane and MA lattice were then progressively brought together by the previously described adsorption/perturbation simulations, with constant ratio pressure control and a pressure of 1 atm via a Nose-Hoover barostat with a period of 20 ps and decay time of 10 ps. The temperature was maintained at 310K using a Langevin thermostat with Langevin friction coefficient reduced from 10 ps^-1^ to 5 ps^-1^ and 1ps^-1^ over periods of 20ns for a full simulation time of 60ns. This approach progressively changes the chemical environment that the MA lattice is exposed to, allowing us to study the structural stability of the lattice and energetic favorability of MA-MA interfaces.

Through the simulations we calculate the distance between neighboring MA trimers as

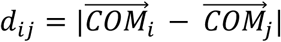

where 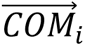 is the center of mass of the i-th MA trimer in the lattice. We identified lattice-breaking events as those where the trimer-trimer distance was greater than the maximum trimer-trimer distance measured from the CryoEM density and WT MA equilibration (*d_ij_* > 55.0Å). Similarity between the trimer-trimer distance distributions was measured using the Jensen-Shannon distance metric [43, 44]. This is a symmetric similarity metric bound between 0 and 1, where two probability distributions will have a distance further from 0 the more dissimilar they are.

We estimated the binding affinity between pairs of neighboring MA trimers by utilizing the molecular mechanics generalized solvent surface area (MM-GBSA) [45, 46, 104] method. MM-GBSA allows the calculation of free binding energies post simulation by calculating interaction energy between two groups of atoms from the molecular mechanics potentials and approximating bulk effects in the long-range electrostatic and Van der Waals interactions with the generalized born and surface area continuum solvation model. MM-PBSA/GBSA has been used recently to estimate protein-protein binding affinities in molecular complexes [47, 105, 106]. We calculate the difference between MA-MA trimer binding affinities of mutant MA compared to WT MA from the MA lattice perturbation simulations using the MMPBSA.py tool [45] available in Amber22 [107], using the igb=5 generalized Born solvation model with 150mM salt concentration.

## Acknowledgments

Research in the Freed laboratory is supported by the Intramural Research Program of the Center for Cancer Research, National Cancer Institute, National Institutes of Health. YH was supported by JSPS Research Fellowship for Japanese Biomedical and Behavioral Researchers at NIH. HV was supported in part by an Intramural AIDS Research Fellowship. The Perilla group is funded by NIH award R01AI178846. This work was supported by the National Institutes of Health grant R21AI184080 and R01AI150560, the UK Wellcome Trust Investigator Award 206422/Z/17/Z, the ERC AdG grant 101021133 and the Chinese Academy of Medical Sciences Oxford Institute core grant CNR00050. We would like to acknowledge our collaborative interactions with the Pittsburgh Center for HIV Protein Interactions (U54AI170791) and the Behavior of HIV in Viral Environments Center (U54AI170855). We acknowledge Diamond Light Source for access and support of the cryoEM facilities at the UK national electron Bio-Imaging Centre (eBIC) (proposal NT29812). Computation was performed at the Oxford Biomedical Research Computing (BMRC) facility supported by the Wellcome Trust Core Award Grant Number 203141/Z/16/Z with additional support from the NIHR Oxford BRC. This work used Delta at NCSA and Stampede3 at TACC through allocation MCB-170096 from the Advanced Cyberinfrastructure Coordination Ecosystem: Services & Support (ACCESS) program, which is supported by National Science Foundation awards #2138259, #2138286, #2138307, #2137603, and #2138296. Support from the University of Delaware CBCB Bioinformatics Data Science Core Facility (RRID:SCR_017696), including use of the BioStore computational resources, was made possible through funding from Delaware INBRE (P20GM103446), NIH Shared Instrumentation Grant (S10OD028725) the State of Delaware, and the Delaware Biotechnology Institute. We thank Drs. Tsutomu Murakami and Kanai Tapan for the technical assistance and suggestions.

## Author contributions

EOF and PZ conceived the project. YH initiated the project, characterized the properties of virus particles carrying WT and mutant MA using biochemical and virological assays. YH purified WT and mutant virus particles for structural studies. LC, YH and YS prepared the cryoEM grids. LC and YZ collected cryoET data. LC performed structural analysis using cryoET and subtomogram averaging. LC developed codes and analyzed MA and CA spatial organization and virion density profile. JRP designed and developed MD methodologies and analyses framework for all simulations. JSR performed molecular dynamics (MD) simulations of MA-lipid interactions for the immature WT and mutant MA lattices as well as mature WT and mutant MA in the presence and absence of membrane. YH and HV conducted MA mutagenesis and infectivity assays for the single amino acid mutants. JRP and JSR analyzed MD simulation results. CA purified WT and mutant proteins, checked by western blotting and imaged them by cryoEM. LC coordinated the project. LC, YH, JSR, PZ, EOF and JRP wrote the manuscript with contributions from all authors.

## Data and materials availability

The immature and mature HIV-1 lattice structures are deposited in the Electron Microscopy Data Bank (EMDB) and Protein Data Bank (PDB) under accession codes: EMD-47842 (https://www.ebi.ac.uk/emdb/EMD-47842) and PDB ID 9EK1 (https://doi.org/10.2210/pdb9ek1/pdb) for the mature MA trimer of wild type, EMD-52051(https://www.ebi.ac.uk/emdb/EMD-52051) for the immature CA hexamer with MA-L20K/E73K/A82T mutations, EMD-52052 (https://www.ebi.ac.uk/emdb/EMD-52052) for the immature CA hexamer of wild type, EMD-52059 (https://www.ebi.ac.uk/emdb/EMD-52059) and PDB ID 9EK2 (https://doi.org/10.2210/pdb9ek2/pdb) for the immature MA trimer with MA-L20K/E73K/A82T mutations, and EMD-52060 (https://www.ebi.ac.uk/emdb/EMD-52060) and PDB ID 9EK3 (https://doi.org/10.2210/pdb9ek3/pdb) for the immature MA trimer of wild type.

## Supplementary Information

**Table S1.** Cryo-EM data acquisition and image processing. *85 out of 171 total particles were randomly selected for STA analysis of the CA.

| Sample | Immature WT |  | Immature L20K/E73K/A82T |  | Mature WT | Mature L20K/E73K/A82T |
| --- | --- | --- | --- | --- | --- | --- |
| Data acquisition |  |  |  |  |  |  |
| Microscope | FEI Titan Krios |  | FEI Titan Krios |  | FEI Titan Krios | FEI Titan Krios |
| Energy-filter slit width (eV) | 10 |  | 10 |  | 10 | 10 |
| Detector | Falcon4 |  | Falcon4 |  | Falcon4 | Falcon4 |
| Pixel size (Å) | 1.50 |  | 1.50 |  | 1.50 | 1.50 |
| Defocus range (microns) | -1.5 to -5.5 |  | -1.5 to -5.5 |  | -1.5 to -5.5 | -1.5 to -5.5 |
| Acquisition scheme | -60°/60°, 3° dose symmetric |  | -60°/60°, 3° dose symmetric |  | -60°/60°, 3° Dose symmetric | -60°/60°, 3° dose symmetric |
| Total Dose (electrons/Å²) | 123 |  | 123 |  | 123 | 123 |
| Frame number | 10 |  | 10 |  | 10 | 10 |
| Tomogram number | 89 |  | 78 |  | 88 | 93 |
| Image processing |  |  |  |  |  |  |
| Virus particles | MA | CA | MA | CA | MA | MA |
|  | 171 | 85/171* | 109 | 109 | 343 | 428 |
| Subtomogram | 18262 | 30782 | 8600 | 30646 | 47770 | Null |
| Symmetry | C3 | C6 | C3 | C6 | C3 | Null |
| Resolution at 0.143 FSC (Å) | 8.0 | 4.9 | 8.3 | 5.2 | 7.3 | Null |
| EMDB accession codes | EMD-52060 | EMD-52052 | EMD-52059 | EMD-52051 | EMD-47842 | Null |

**Table S2.** HIV-1 asymmetric flat lipid bilayer lipid composition. For each lipid species present in the membrane (specified by their CHARMM topology name), we list the total number of molecules in the membrane as well as the molar fraction and the fraction of molecules present in the intravirion and extravirion leaflet. The intravirion leaflet is rich in PIPs, phosphatidylethalonamine (PE) and phosphatidylserine (PS) while the outer leaflet is enriched in sphingomyelins (SM), ceramides (CER), glycerides (GL) and phosphatidylcholine (PC). The intravirion and extravirion leaflets contain 1797 and 1964 lipid molecules, respectively, for a total of 3761 lipids in the bilayer membrane.

| Lipid (CHARMM) | No.Molecules (Total) | Mol.fraction (%) | Intravirio ratio | Extravirion ratio | No.Molecules (Intravirion) | No.Molecules (Extravirion) |
| --- | --- | --- | --- | --- | --- | --- |
| CER160 | 8 | 0.21 | 0.12 | 0.88 | 1 | 7 |
| CER180 | 3 | 0.08 | 0.12 | 0.88 | 1 | 2 |
| CHL1 | 1260 | 33.50 | 0.47 | 0.53 | 556 | 704 |
| DLPC | 83 | 2.21 | 0.2 | 0.8 | 15 | 68 |
| DOPC | 48 | 1.28 | 0.2 | 0.8 | 8 | 40 |
| DPPC | 109 | 2.90 | 0.28 | 0.72 | 25 | 84 |
| NSM | 129 | 3.43 | 0.12 | 0.88 | 10 | 119 |
| PAPE | 214 | 5.69 | 0.75 | 0.25 | 155 | 59 |
| PAPS | 78 | 2.07 | 0.81 | 0.19 | 62 | 16 |
| PDOPC | 15 | 0.40 | 0.18 | 0.82 | 3 | 12 |
| PDOPE | 217 | 5.77 | 0.92 | 0.08 | 197 | 20 |
| PEPE | 27 | 0.72 | 0.9 | 0.1 | 24 | 3 |
| PEPS | 127 | 3.38 | 0.75 | 0.25 | 93 | 34 |
| PLPE | 140 | 3.72 | 0.85 | 0.15 | 117 | 23 |
| PLPS | 84 | 2.23 | 0.76 | 0.24 | 62 | 22 |
| POPC | 141 | 3.75 | 0.16 | 0.84 | 19 | 122 |
| POPE | 20 | 0.53 | 0.75 | 0.25 | 15 | 5 |
| POPS | 26 | 0.69 | 0.74 | 0.26 | 18 | 8 |
| SDPC | 9 | 0.24 | 0.2 | 0.8 | 1 | 8 |
| SDPE | 146 | 3.88 | 0.88 | 0.12 | 126 | 20 |
| SDPS | 190 | 5.05 | 0.73 | 0.27 | 134 | 56 |
| SSM | 529 | 14.07 | 0.12 | 0.88 | 38 | 491 |
| SAPI | 42 | 1.12 | 0.85 | 0.15 | 35 | 7 |
| SAPI14 | 19 | 0.51 | 0.85 | 0.15 | 16 | 3 |
| SAPI24 | 68 | 1.81 | 0.85 | 0.15 | 57 | 11 |
| SAPI34 | 1 | 0.03 | 0.85 | 0.15 | 1 | 0 |
| POGL | 28 | 0.74 | 0.34 | 0.66 | 8 | 20 |
| Total | 3761 |  |  |  | 1797 | 1964 |

**Table S3.**
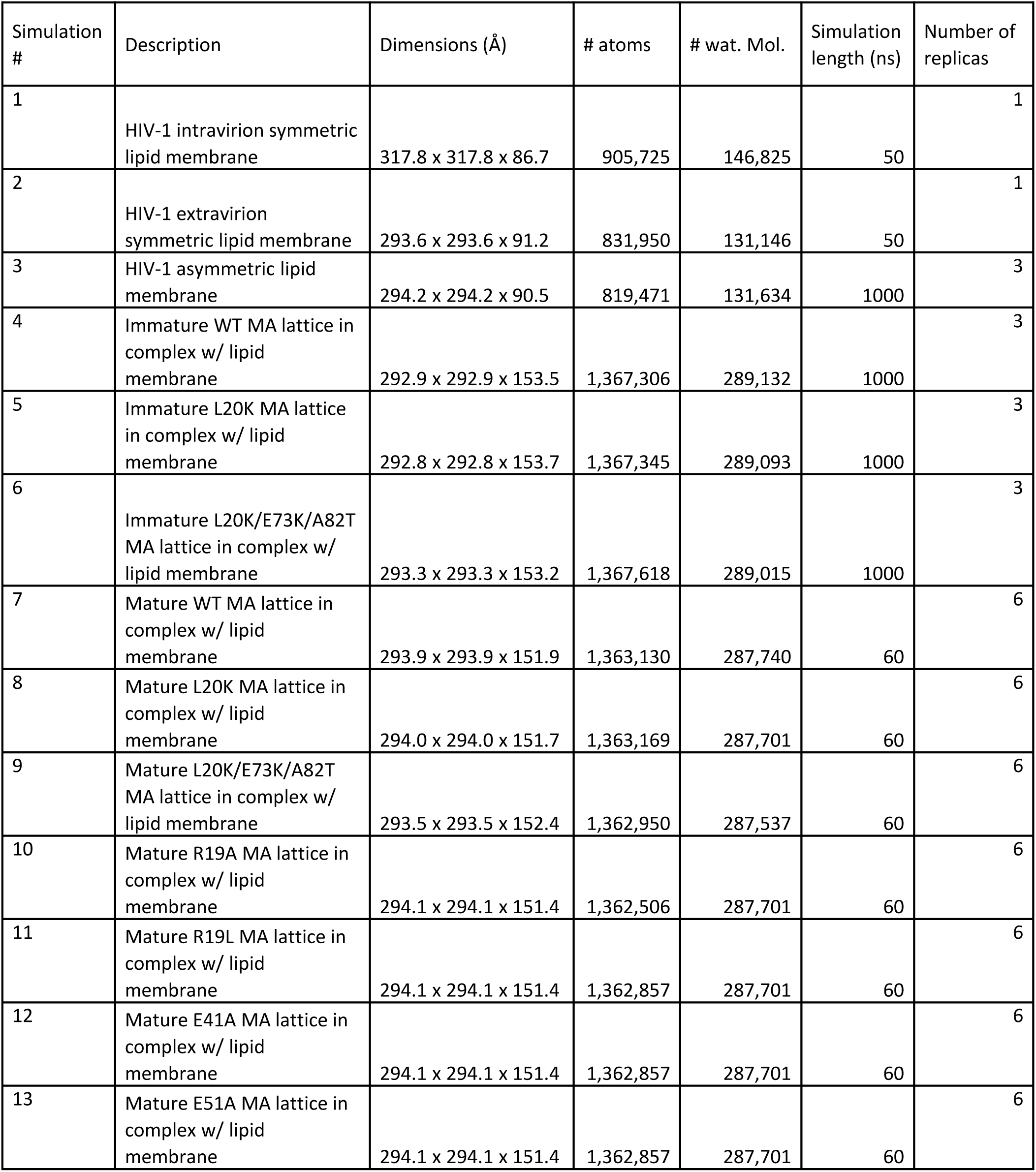
Lipid membrane and membrane in complex with MA systems built for MD simulations. The symmetric lipid bilayers (#1,#2) which were prepared using CHARMM-GUI [76] and equilibrated shortly before using the lipid coordinates as templates to build the asymmetric lipid membrane (#3). The apo HIV-1 asymmetric lipid membrane and in complex with the immature MA lattices (WT, L20K and L20K/E73K/A82T) were simulated for 1 μs (#4-#6) before calculating lateral lipid displacement, lipid headgroup occupancy and protein-lipid contact occupancies. Mature MA lattice perturbation simulations (#7-#13) via adsorption to the lipid membrane were performed for 60 ns varying the Langevincoupling constant from 10 ps^-1^ to 5ps^-1^ and 1 ps^-1^ over periods of 20 ns allowing the MA and membrane to progressively come together and probing the MA-MA intertrimer and intratrimer interactions and assess the MA lattice structural stability. All simulations were performed on NPT ensembles with constant ratio control.

| Simulation # | Description | Dimensions (Å) | # atoms | # wat. Mol. | Simulation length (ns) | Number of replicas |
| --- | --- | --- | --- | --- | --- | --- |
| 1 | HIV-1 intravirion symmetric lipid membrane | 317.8 x 317.8 x 86.7 | 905,725 | 146,825 | 50 | 1 |
| 2 | HIV-1 extravirion symmetric lipid membrane | 293.6 x 293.6 x 91.2 | 831,950 | 131,146 | 50 | 1 |
| 3 | HIV-1 asymmetric lipid membrane | 294.2 x 294.2 x 90.5 | 819,471 | 131,634 | 1000 | 3 |
| 4 | Immature WT MA lattice in complex w/ lipid membrane | 292.9 x 292.9 x 153.5 | 1,367,306 | 289,132 | 1000 | 3 |
| 5 | Immature L20K MA lattice in complex w/ lipid membrane | 292.8 x 292.8 x 153.7 | 1,367,345 | 289,093 | 1000 | 3 |
| 6 | Immature L20K/E73K/A82T MA lattice in complex w/ lipid membrane | 293.3 x 293.3 x 153.2 | 1,367,618 | 289,015 | 1000 | 3 |
| 7 | Mature WT MA lattice in complex w/ lipid membrane | 293.9 x 293.9 x 151.9 | 1,363,130 | 287,740 | 60 | 6 |
| 8 | Mature L20K MA lattice in complex w/ lipid membrane | 294.0 x 294.0 x 151.7 | 1,363,169 | 287,701 | 60 | 6 |
| 9 | Mature L20K/E73K/A82T MA lattice in complex w/ lipid membrane | 293.5 x 293.5 x 152.4 | 1,362,950 | 287,537 | 60 | 6 |
| 10 | Mature R19A MA lattice in complex w/ lipid membrane | 294.1 x 294.1 x 151.4 | 1,362,506 | 287,701 | 60 | 6 |
| 11 | Mature R19L MA lattice in complex w/ lipid membrane | 294.1 x 294.1 x 151.4 | 1,362,857 | 287,701 | 60 | 6 |
| 12 | Mature E41A MA lattice in complex w/ lipid membrane | 294.1 x 294.1 x 151.4 | 1,362,857 | 287,701 | 60 | 6 |
| 13 | Mature E51A MA lattice in complex w/ lipid membrane | 294.1 x 294.1 x 151.4 | 1,362,857 | 287,701 | 60 | 6 |

**Table S4.** Cryo-ET data acquisition and image processing for L20K.

| MA | Immature L20K | Mature L20K |
| --- | --- | --- |
| Data acquisition |  |  |
| Microscope | FEI Titan Krios | FEI Titan Krios |
| Energy-filter (eV) | 20 | 20 |
| Detector | Falcon4 | Falcon4 |
| Pixel size (Å) | 1.50 | 1.50 |
| Defocus range (microns) | -1.5 to -5.5 | -1.5 to -5.5 |
| Acquisition scheme | -60°/60°, 3°, dose symmetric | -60°/60°, 3°, dose symmetric |
| Total Dose (electrons/Å²) | 123 | ~123 |
| Frame number | 10 | 10 |
| Tomogram number<br>(processed/total collected) | 16 | 11 |
| Image processing |  |  |
| Virus particles | 123 | 97 |
| Wild type MA lattice<br>template matching analysis | Yes | Yes |

**Figure S1.**
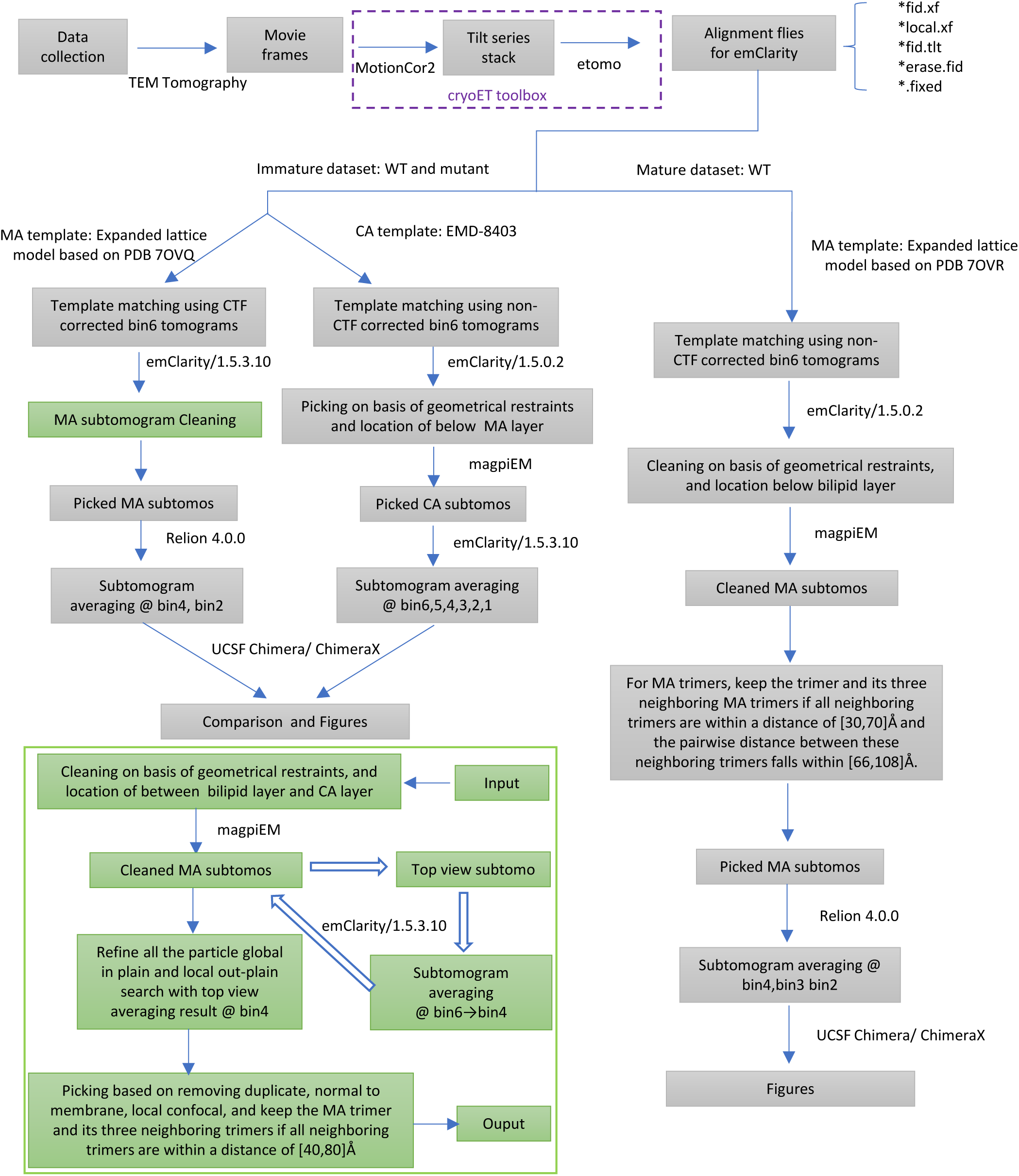
Workflow for data processing. TEM Tomography 5 was utilized to capture Falcon4 tilt-series. On-the-fly pre-processing was conducted via the CryoET Toolbox (https://github.com/ffyr2w/cet_toolbox). This process included frame motion correction via MotionCor2 [63] and fiducial marker-based alignment through etomo [64]. The resulting alignment files were then transferred to emClarity for template matching [39, 65]. Template matching outcomes was cleaned for both CA and MA based on geometric constraints with in-house-developed script magpiEM (https://github.com/fnight128/MagpiEM). The cleaned subtomogram selection dataset was randomly split into two subsets for parallel processing. The gold-standard Fourier shell correlation was computed from separate even and odd halves. For immature CA, we selected particles from 6x binned, non-CTF-corrected tomograms through emClarity/1.5.0.2 template matching, employing a 28 Å low-pass filter on the EMD-8403 template [66]. Hexamer CA’s iterative alignment was conducted from 6x to 1x binning using emClarity/1.5.3.10, maintaining C6 symmetry throughout. The immature MA analysis began with the PDB 7OVQ [18] template to create a 400Å-wide lattice map for template matching on 6x binned, CTF-corrected tomograms using emClarity/1.5.3.10, also with a 28 Å low-pass filter. The cleaning was undertaken by geometric constraints, focusing on the inter layer between the bilipid and CA layers with magpiEM. Due to false positive peaks induced by the missing wedge from side views, top view subtomograms were selected firstly, restricting the choice to those peaks with a tilt angle of less than 45 degrees relative to the beam direction. Subsequent averaging for top view subtomograms was performed from bin6 to bin4. We refined all subtomograms globally in-plane and locally out-of-plane, ensuring top view coordinates aligned with those refined at bin4. MA subtomograms were picked by removing duplicates and satisfying local lattice constrain, readying them for subtomogram averaging in Relion 4.0.0 with global refinement at bin4 and local refinement at bin2 with C3 being enforced [38]. For mature wild-type MA, a mature expanded 400Å-wide lattice model based on PDB 7OVR was used as the template [18]. Template matching is operated on 6x binned, non-CTF-corrected tomograms in emClarity/1.5.0.2, using a 28 Å low-pass filter. The matching result was cleaned based on geometric restraints and location below the bilipid layer with magpiEM. Further cleaning of trimeric MA was guided by criteria that retained a trimer and its three neighboring trimers if all neighboring trimers fell within a distance of [30,70] Å and the pairwise distances between these neighboring trimers ranged from [66,108] Å. This process removed unreliable particle peak, yielding a dataset of cleaned subtomograms. These subtomograms were further processed using Relion 4.0.0, where subtomogram averaging was initially performed at bin4 with global refinement, then followed by local refinement at bin3 and bin2, enforcing C3 symmetry throughout. ChimeraX [68] and ArtiaX [67] were used for visualization and analysis of the density maps, supplemented by false discovery rate analyses [69].

**Figure S2.**
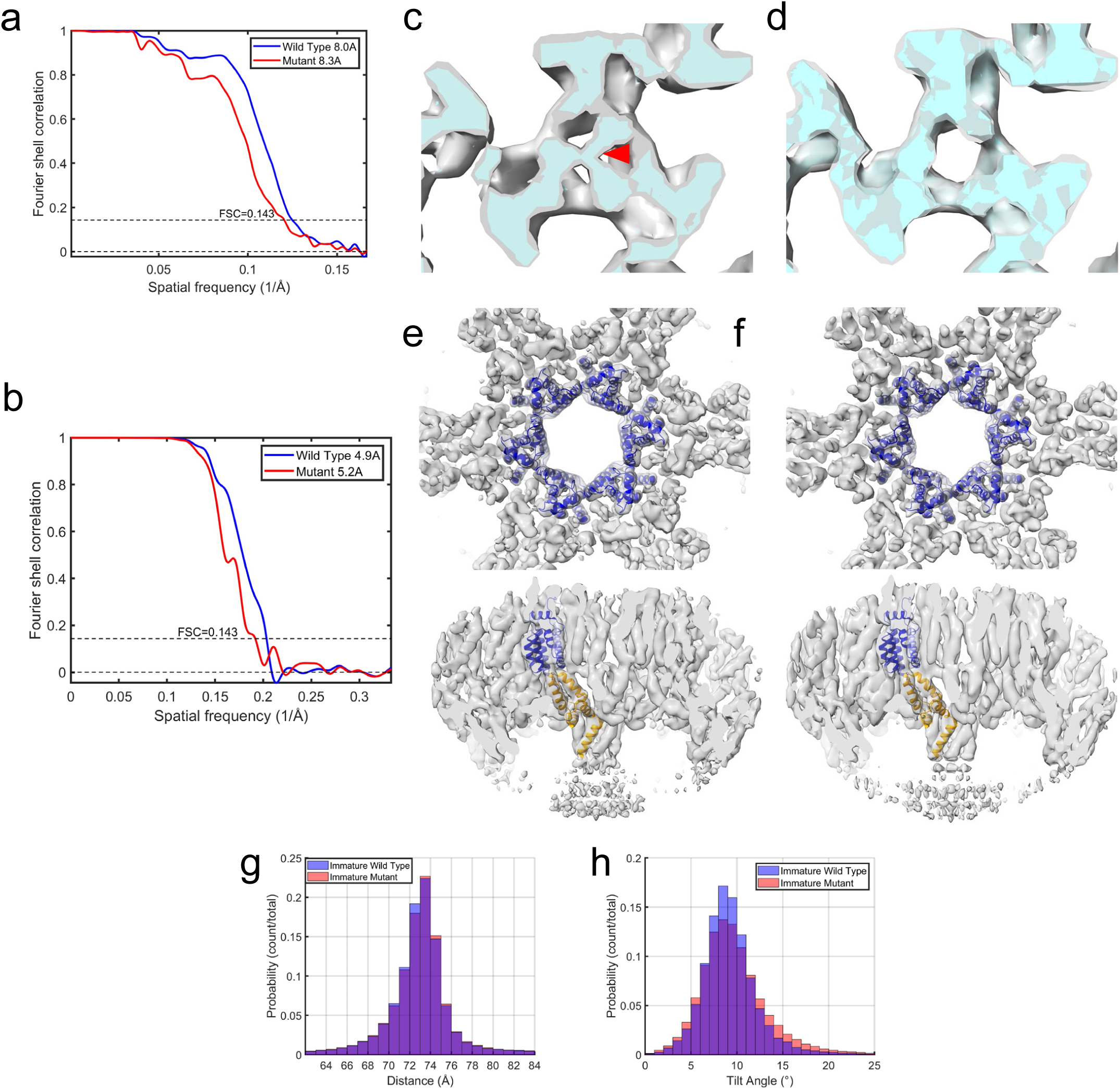
Structures of immature MA and CA. a-b) Fourier shell correlation (FSC) plots for the MA trimer (a) and CA hexamer (b) from both immature wild type and mutant, with resolutions indicated at the FSC value of 0.143. c-d) Final refined maps for MA wild type (c) and mutant (d) with C1 symmetry, displaying distinct central differences. The top view of the confidence map, calculated at a 1% false discovery rate [69], is shown in cyan, superimposed on the gray MA trimer map. (e-f) The difference map from Fig. 3c is overlaid on the wild-type MA trimer map. e-f) CA hexamer maps for the wild type (e) and mutant (f), displayed in top and side views, are superimposed with the model (PDB 5L93) [110]. The N-terminal domain (NTD) and C-terminal domain (CTD) of CA are colored blue and orange, respectively. (g-h) Comparisons of distances and tilt angles between hexamer-hexamer pairings in tomographic reconstructions reveal that the CA-CA distance for the WT averages 72.9 Å with a standard deviation of 2.9 Å, compared to the mutant’s mean of 73.0 Å and standard deviation of 2.9 Å. Tilt angles for the WT and mutant are 9.1° (SD 2.7°) and 9.4° (SD 3.3°) respectively.

**Figure S3.**
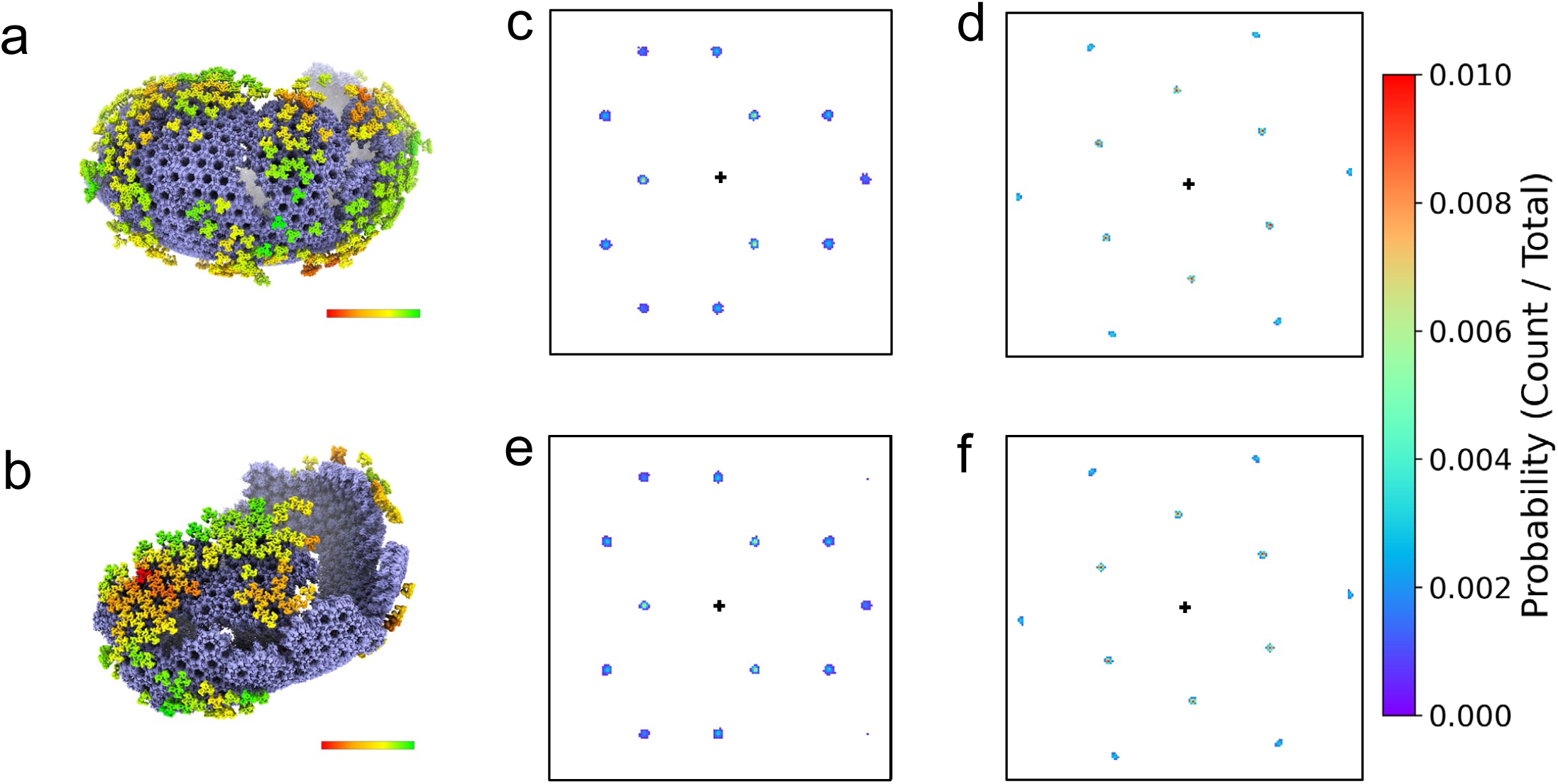
lattice regularity comparison of MA and CA in immature wild type and L20K/E73K/A82T mutant VLPs. a-b) Lattice maps of MA trimer (colored) and CA hexamer (blue) from an example tomogram of WT (a) and mutant (b) VLPs. MA trimers are colored by cross correlation values with the average MA trimer map, from red (lower) to green (higher), with ranges of 1.501e6 to 1.512e6 for wild type and 1.459e6 to 1.471e6 for mutant. c-f) Localization plots for MA trimers and CA hexamers illustrate the positions of neighboring units around a central trimer or hexamer (black cross). In WT VLPs, there are 18,262 MA trimers (c) and 30,782 CA hexamers (d). In mutant VLPs, there are 8,600 MA trimers (e) and 30,646 CA hexamers (f). The color gradient in the heatmaps, from blue to red, indicates the increasing probability for the occurrence of neighbouring trimers or hexamers as per the color key.

**Figure S4.**
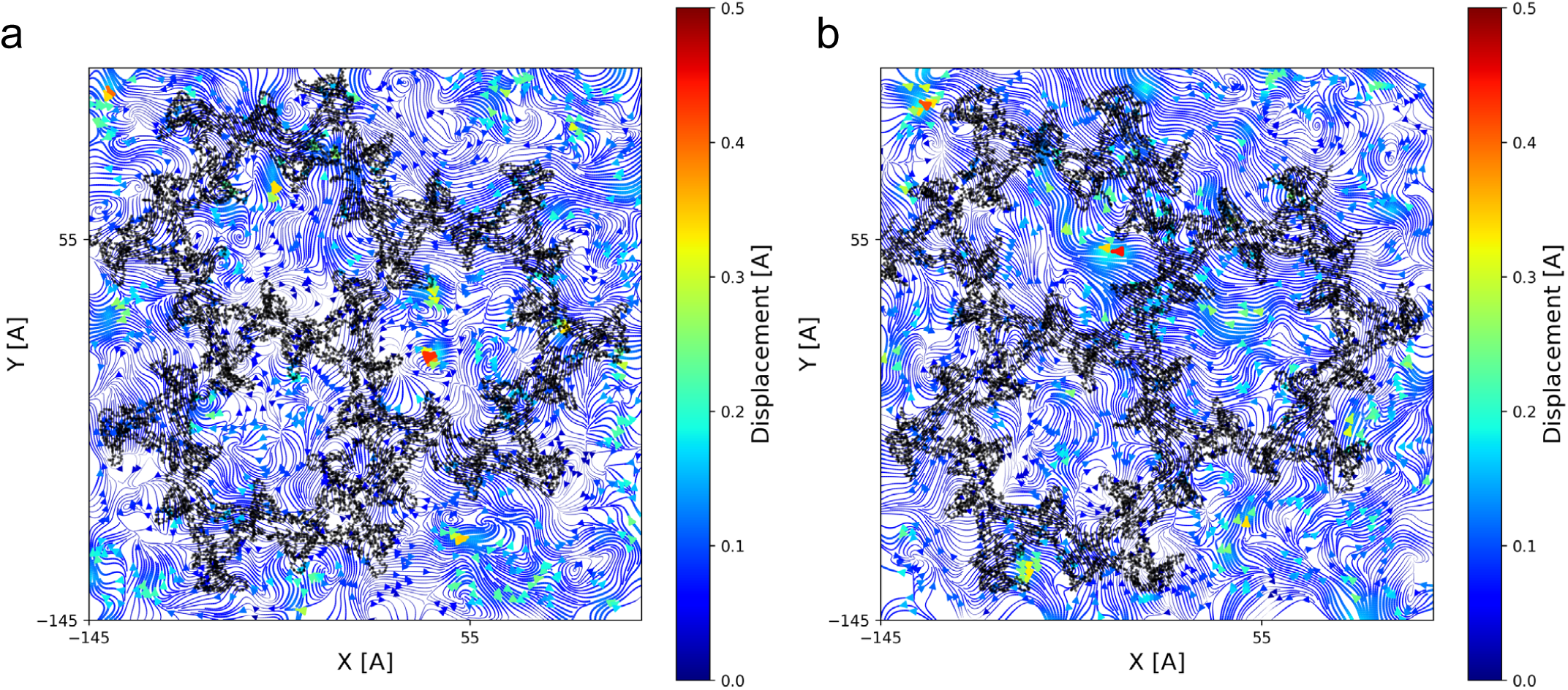
Instantaneous lateral lipid displacement visualization. a-b) Streamline visualization of the instantaneous lateral lipid displacement of lipid headgroups in the intravirion leaflet in the presence of the immature WT MA lattice (a) or the immature L20K/E73K/A82T MA lattice. MA lattice position is colored in black.

**Figure S5.**
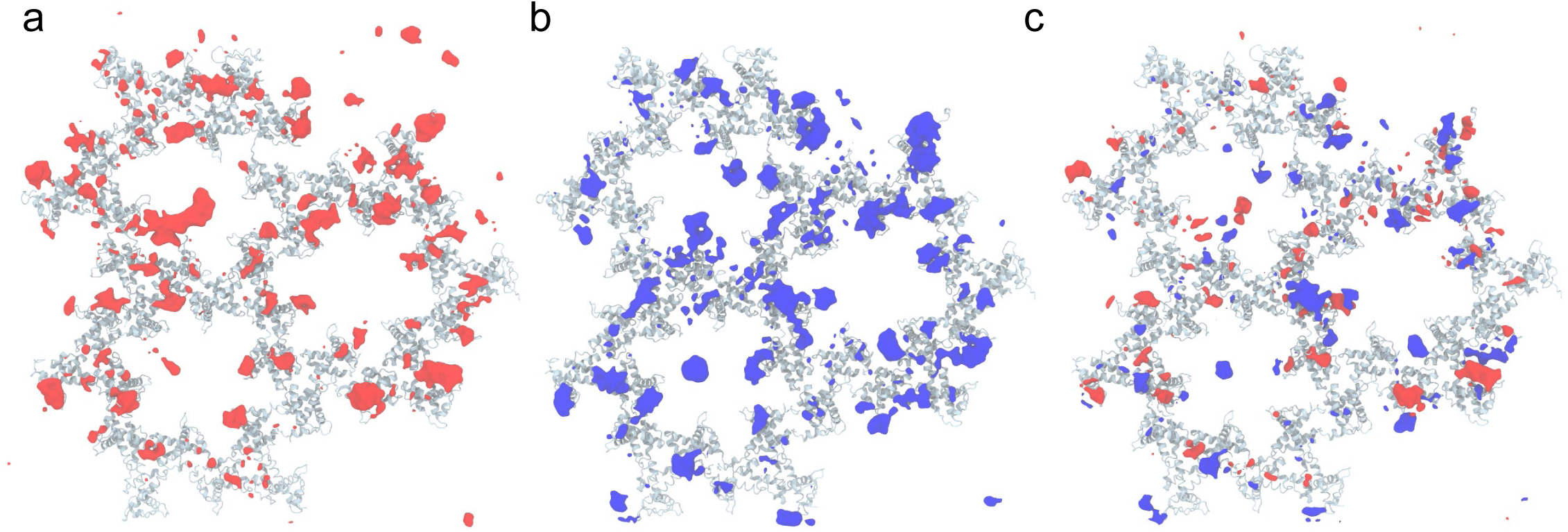
Lipid headgroup occupancies in presence of immature MA. a-b) Occupancy maps for lipid headgroups through 1 μs of MD simulation for the WT MA (a) or L20K/E73K/A82T MA (b) at isovalue >25%. c) Lipid occupancy difference map, calculated as L20K/E73K/A82T MA – WT MA. Red volumes represent regions present in the WT MA lipid occupancy map and not the L20K/E73K/A82T MA lipid occupancy map. Blue volumes represent regions present in the L20K/E73K/A82T MA lipid occupancy map and not the WT MA lipid occupancy map. Immature MA lattice is shown in light blue in all panels.

**Figure S6.**
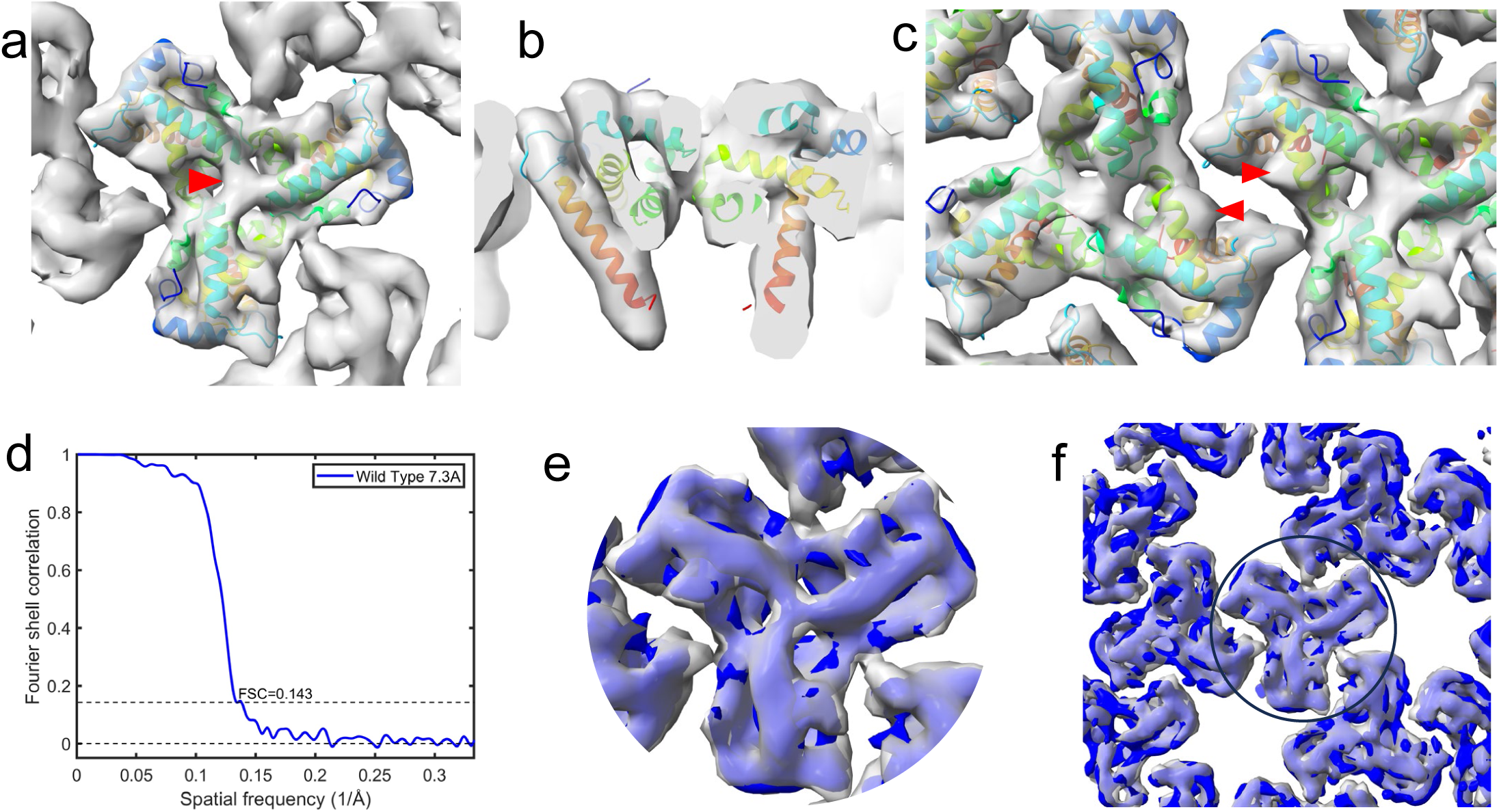
Subtomogram averaging and FSC of mature wild type MA lattice. a-b) The subtomogram averaging map of the wild type MA trimer is represented in gray isosurface, viewed from the top towards the center of the virus and from the side perpendicular to the membrane. The molecular model of 7OVR [18] is superimposed as a rigid body, color-coded from blue at the N-terminus to red at the C-terminus. c) Density at the PI(4,5)P2 binding site is highlighted by red arrowheads. d) FSC curve assessing the resolution of the MA lattice structure, indicating resolution values at the 0.143 threshold. e-f) The mature MA trimer (e) and its lattice (f) map are overlaid with the the previously published EMD-13088 map [18]. Our MA density is shown in grey and the density of EMD-13088 is in blue.

**Figure S7.**
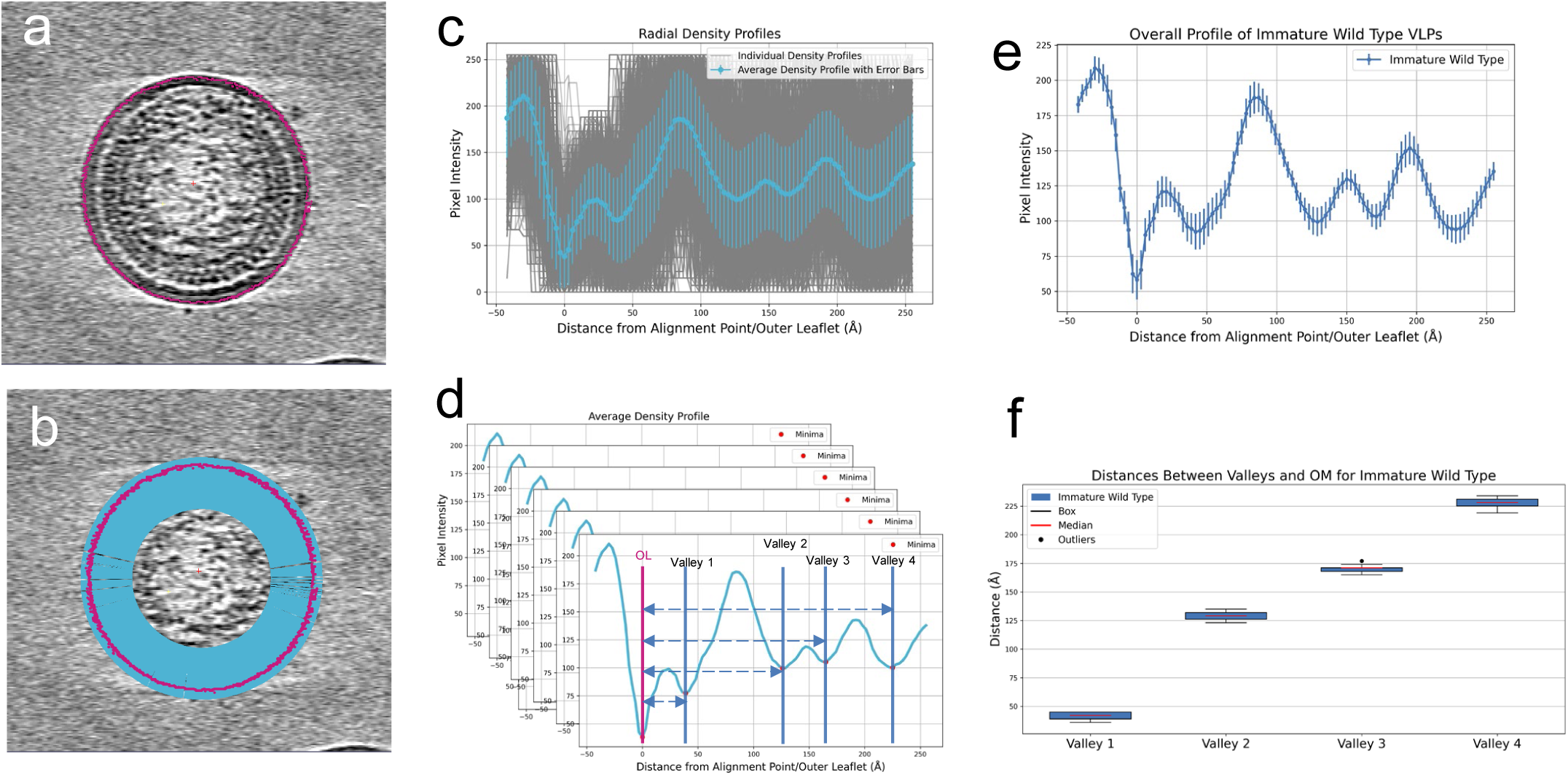
Profile analysis method: Outer membrane-aligned profiling and averaging. a) Identify the position of the outer leaflet (OL). b) Generate the density profiles perpendicular to OL. c) Compute the average across all profiles aligned based on the OL position. d) Accumulate a sufficient number of VLP average profiles for robust analysis. e) Calculate the overall average profile, presented with error bars. f) Plot a box plot to analyze the distances between various valleys and the OL.

**Figure S8.**
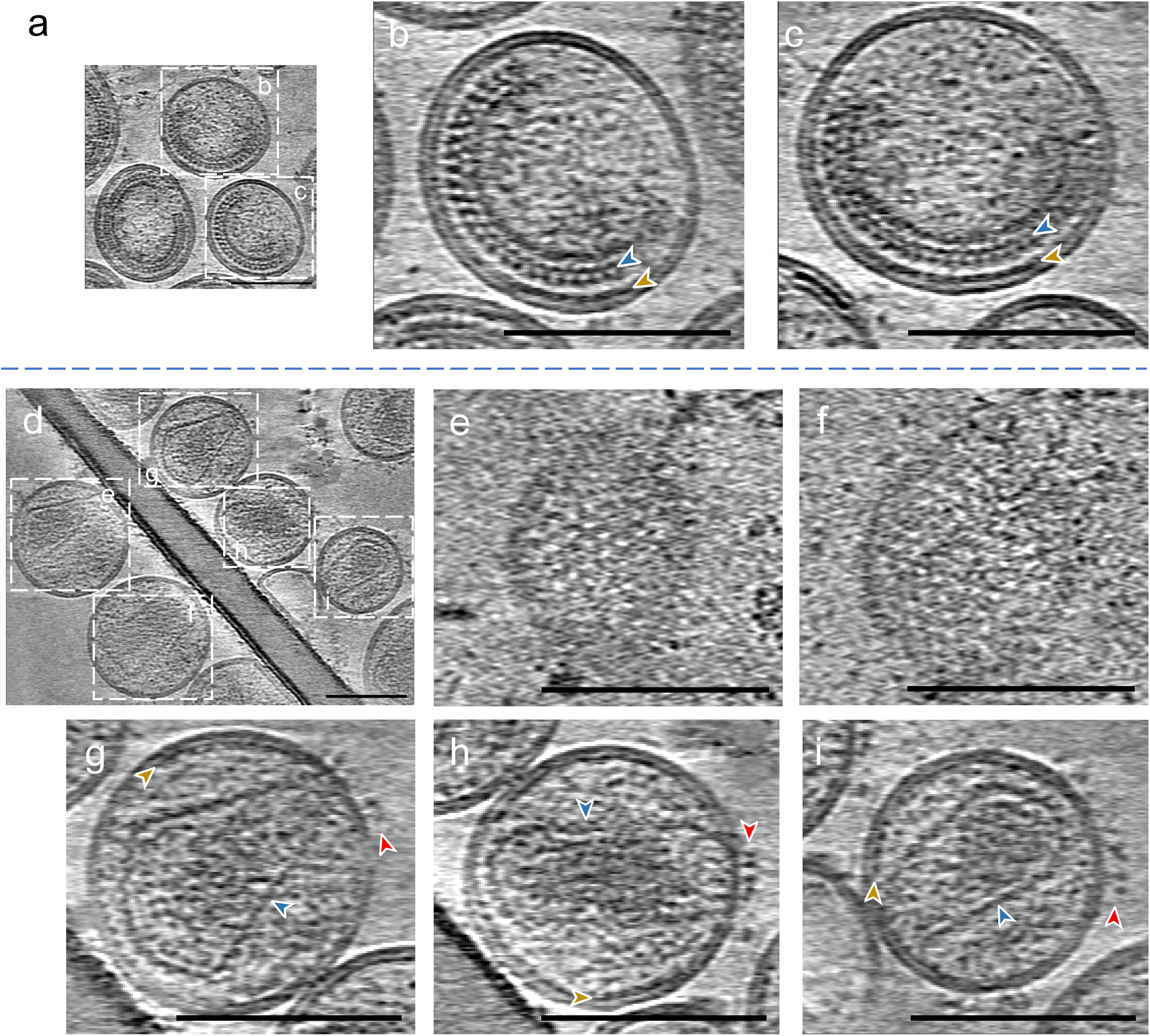
Tomograms of immature and mature MA L20K viruses. a) Representative central slice of immature L20K particles, enhanced by summing 10 adjacent slices. b-c) Detailed views of the central sections from the boxed regions b and c in panel a, similarly enhanced, exhibit the lattice pattern associated with the inner membrane, brown arrowheads mark the positions of the MA lattice, blue arrowheads indicate the CA lattice. d) Representative central slice of mature L20K particles, enhanced by summing 10 neighboring slices. e-f) Magnified top view of slices near the inner membrane surface from boxed areas e and f in panel d, showing lattice patterns. g-i) Magnified central slices from boxed areas g to i in panel d, also enhanced by summing 10 neighboring slices, showing the lattice pattern beneath the inner membrane. Brown arrowheads indicate the MA lattice, blue arrowheads point to the capsid, and red arrowheads highlight the Env glycoproteins. Scale bar: 100nm.

**Figure S9.**
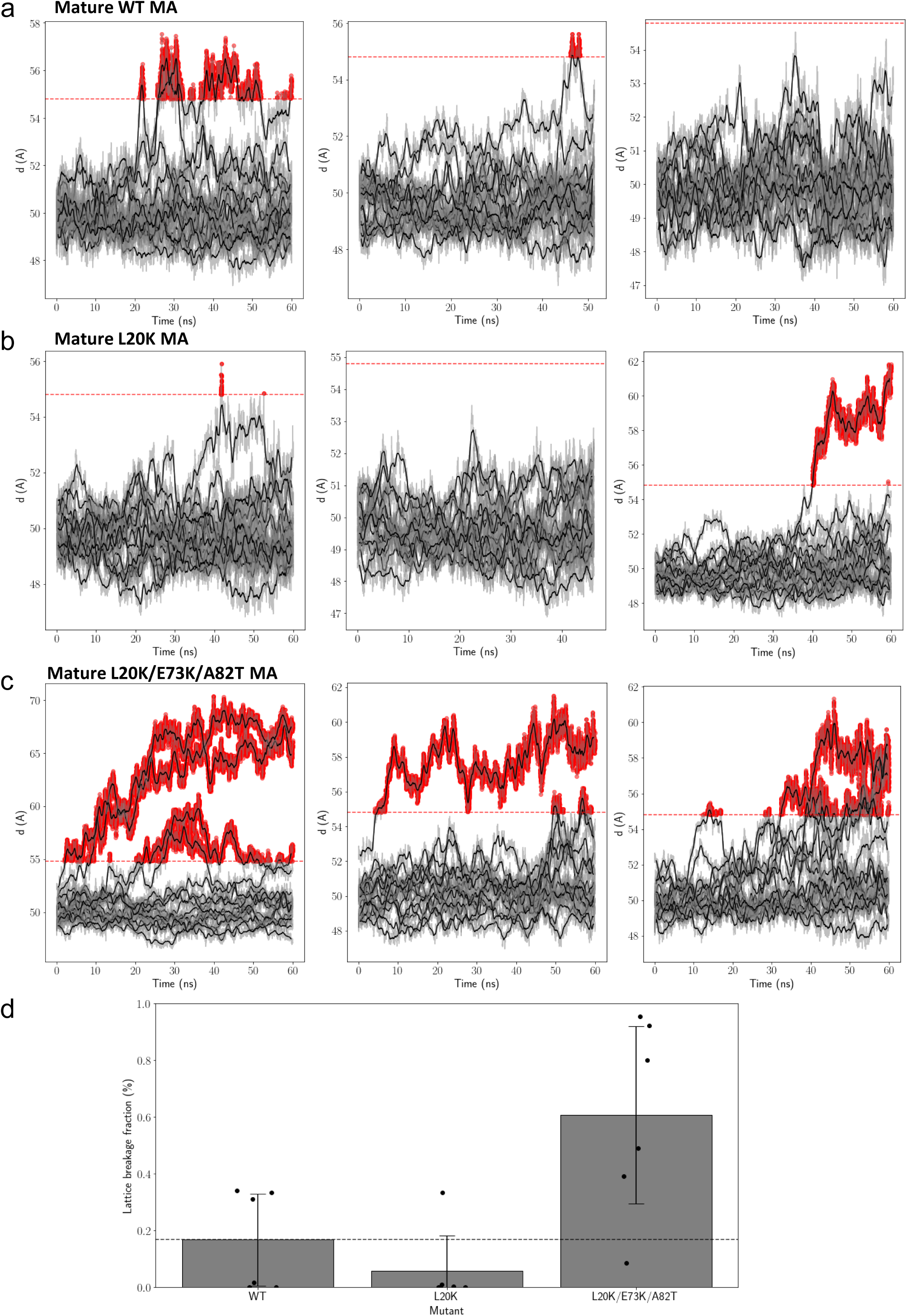
MA trimer-trimer distance traces from perturbation simulations. a) mature WT MA lattice b) mature L20K MA lattice and c) mature L20K/E73K/A82T MA lattice. Trimers separating in lattice breaking event are colored in red. 3 out of 6 replicas performed are shown. d) Fraction of lattice breakage events though the 6 replicas.

**Figure S10.**
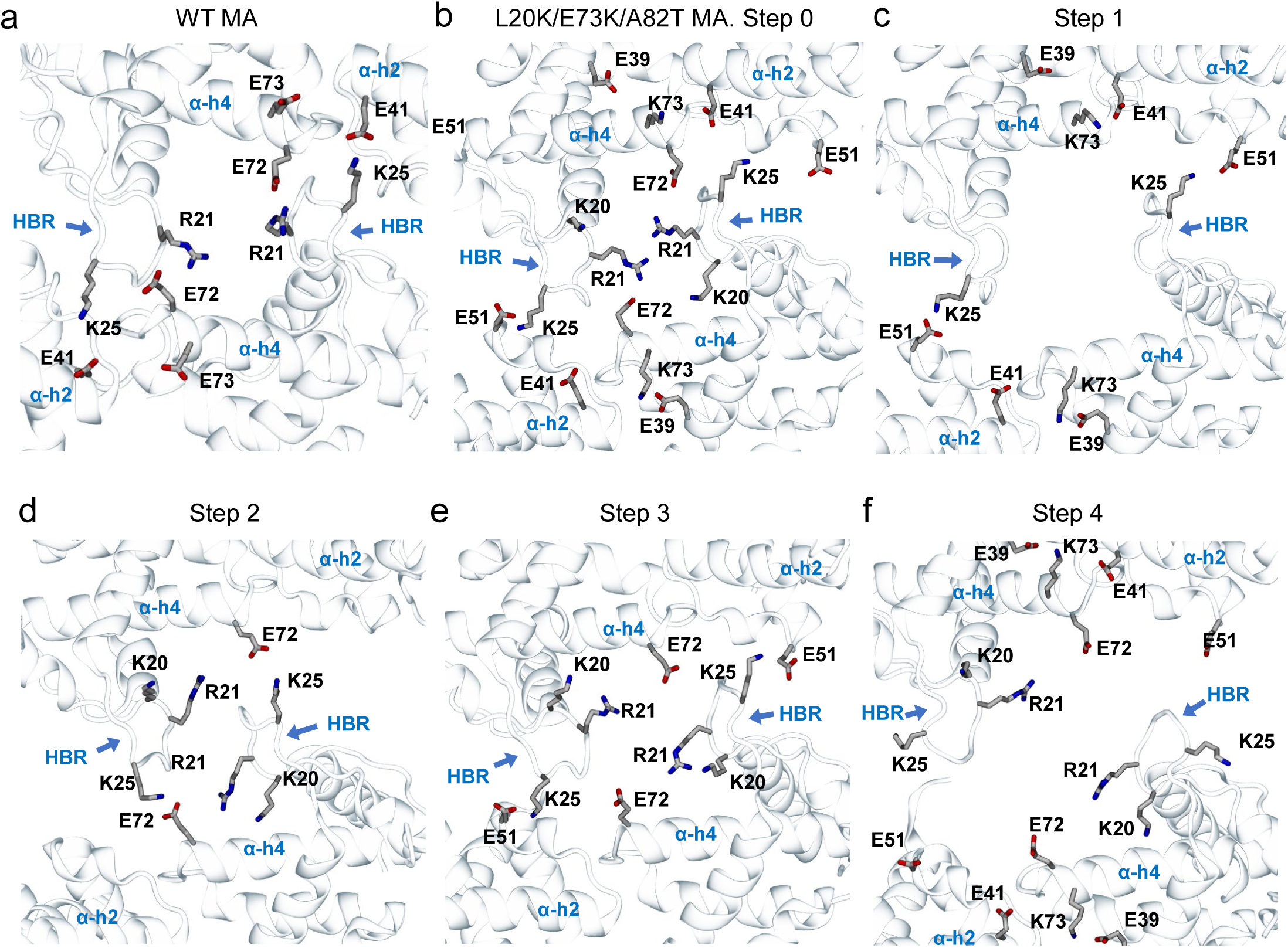
Allosteric disruption of the mature MA inter-trimer interface by the L20K/E73K mutations. a) Reference salt-bridge network at the MA inter-trimer interface for mature WT MA. b-f) Stepwise process for disruption of the MA inter-trimer salt-bridge network. b) E73K interacts with E39 and E41, impeding the salt bridge formation between K25 and E41. c) K25 instead attempts to form salt bridge interactions with E51 and E72. d) E72 is now attracted by positively charged L20K and interacts with R21 in the same monomer. e) R25 forms the only inter-trimer interaction with E51 while other salt bridges are formed intra-trimer. f) Small displacements of the monomers break R51-E51 and the dimer interface is disrupted.

**Figure S11.**
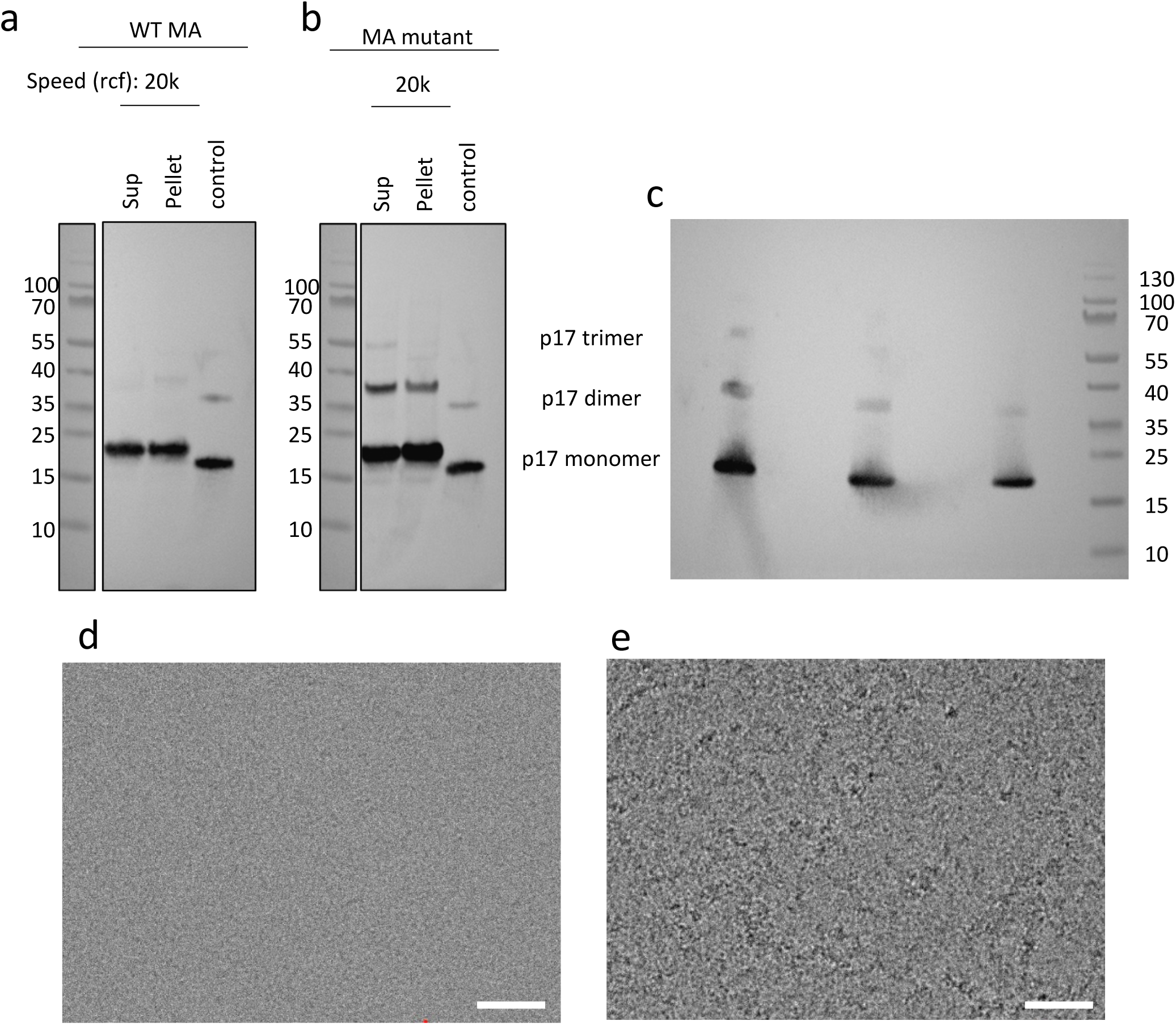
Purification of WT and L20K/E73K/A82T MA. a-b) Western blot analysis of cell lysates expressing N-terminal 8-histidine tagged WT and L20K/E73K/A82T mutant MA using a polyclonal anti-Gag p17 antibody (PAB1178) at a 1:2500 dilution. Sup: Supernatant, Control: Recombinant MA mutant protein. A total of 10 µg of protein was loaded per well. c) SDS-PAGE gel of purified L20K/E73K/A82T MA, with 1, 0.5 and 0.25 μg protein loaded. d-e) Transmission electron micrographs of WT MA (d) and L20K/E73K/A82T MA mutant proteins (e). Scale bar 50 nm.

